# R2HaPpY: Rapid-robust phosphotyrosine peptide enrichment using HaloTag-Src SH2 pY superbinder

**DOI:** 10.1101/2025.05.14.653984

**Authors:** Alexis Chang, Ricard A. Rodriguez-Mias, Matthew D. Berg, Sophie Moggridge, Judit Villén

## Abstract

Phosphotyrosine signaling plays a critical role in many biological processes, from cell proliferation to immune response. Despite its importance, systems-level analysis of phosphotyrosine signaling remains a challenge due to costly enrichment reagents and labor-intensive protocols. We previously established an automated phosphotyrosine enrichment method for preparing 96 samples in parallel. Here, we further optimize this method by fusing an SH2 phosphotyrosine superbinder to the HaloTag protein. This allows simple and cost-effective preparation of enrichment beads directly from bacterial lysate, expediting reagent preparation from days to hours. Additionally, our new reagent binds phosphotyrosine peptides at higher efficiency than other enrichment reagents. Using this reagent, we detect and quantify 1,651 unique phosphotyrosine sites from EGF stimulated HeLa cells using only ∼1 mg of input peptides per replicate. These include 878 regulated pY sites, many of which are low abundance and not previously detected or annotated as EGF-responsive. This streamlined and sensitive method facilitates comprehensive, quantitative mapping of tyrosine phosphorylation dynamics, enabling broader integration of phosphotyrosine signaling into multiomic and network-level models across diverse biological systems and disease states.

## Introduction

Cells respond to external cues via highly interconnected signaling networks^1–7^. Phosphotyrosine (pY) signaling is an important component of intercellular communication systems that regulates many multicellular processes such as coordinated cell proliferation, differentiation, and migration. Phosphotyrosine signaling originates at receptor tyrosine kinases and is transduced through receptor autophosphorylation followed by recruitment and activation of other signaling proteins. Dysregulation of pY signaling is involved in many diseases, especially cancer^8,9^ and the development of tyrosine kinase inhibitors as targeted cancer therapeutics has been highly successful^10^. Systems level measurements of the phosphoproteome have begun to capture how signaling networks are organized and their temporal dynamics in response to perturbation^1,7,11–21^. Deeper understanding of tyrosine signaling, including regulatory dependencies and network connectivity will further advance drug development and illuminate key mechanisms in cellular function. Despite its importance, comprehensive measurement of tyrosine phosphorylation is difficult, due to its low relative abundance and rapid removal by protein tyrosine phosphatases^15^. As a result, global phosphoproteomic enrichment methods that use immobilized metal cations or metal oxides^22–29^ preferentially capture the higher abundance phosphoserine (pS) and phosphothreonine (pT)-containing peptides. This leads to relatively few identified pY-containing peptides.

To improve study of pY signaling, researchers have turned to pY-specific enrichment, typically using pY-specific antibodies^11–13,15,20,30–39^. While effective, pY antibodies are expensive, making large scale studies cost-prohibitive^40^. Not surprisingly, the scaling of pY enrichment to systems-level studies has lagged behind global phosphopeptide studies^1,29,41–47^.

To circumvent the high cost of antibodies, Src Homology 2 (SH2) protein domains, which naturally bind pY^48–54^, have been engineered for high-affinity pY-peptide binding and used as enrichment reagents^40,55–61^. Engineered Src SH2 domains, termed pY superbinder Src SH2s (sSrc), can enrich pY-containing peptides comparably or better than antibodies^40,62–65^ and can be recombinantly expressed and purified from *Escherichia coli*, reducing their production cost relative to antibodies^40,61–67^.

Efforts to study pY modifications at similar ease and affordability as pS or pT have resulted in the development of high throughput and automated pY superbinder-based peptide enrichments^56,65,66^. In 2023, we introduced R2-pY, an automated and high throughput pY enrichment protocol using sSrc immobilized to magnetic beads via lysine reactive N-hydroxysuccinimide (NHS) chemistry and a magnetic particle processing robot^65^; and demonstrated it as a cost-effective, reproducible and efficient approach for enriching pY peptides from up to 96 samples in parallel. Despite the reproducibility and quantitative scalability, R2pY is still burdened by intensive sSrc bead preparation, involving affinity purification of sSrc protein, buffer exchange, and multi-step conjugation; which hinders wide adoption of the method. Similar approaches are used by other groups to prepare the sSrc reagent for both low and high throughput pY enrichments^40,56,57,60–64,66,68,69^.

To overcome these limitations and improve the accessibility of high-throughput sSrc pY-peptide enrichments, we aimed to combine the purification and conjugation of the sSrc into a single step that is easy to perform, bypassing the need for protein chromatography or buffer exchange setups. To achieve this, we fused the sSrc to the HaloTag^70^ protein to enable one-step covalent conjugation to magnetic beads directly from crude *E. coli* lysate. Covalent bond formation between the sSrc- HaloTag fusion protein and the chloroalkane-functionalized magnetic beads is highly specific, rapid in physiological conditions and essentially irreversible^70^. Here, we demonstrate that the HaloTag- sSrc magnetic bead preparation coupled with our automated and high-throughput pY peptide enrichment protocol is simpler and offers unprecedented pY site measurement depth. The method performs well over a range of peptide input amounts and concentrations, and the conjugated beads are stable for at least four months. Importantly, the enrichment beads are 10 times cheaper than pY antibody beads, facilitating cost-effective application in large scale pY studies. We call the improved method R2HaPpY (rapid-robotic <u>Ha</u>lo-sSrc <u>p</u>eptide-level <u>p</u>hosphot<u>y</u>rosine enrichment).

Using this improved method, we profile the signaling response of HeLa cells to epidermal growth factor (EGF) stimulation at 5 distinct time points using only ∼1 mg of protein input per replicate. We identify 1,651 total pY sites, 878 of which significantly change between time points and uncover novel signaling events in the context of the well-studied epidermal growth factor stimulation. Our method not only recapitulates previously observed pY site regulation, but also reproducibly uncovers 70 new EGF-responsive pY sites not before annotated in PhosphoSitePlus^71,72^, many of which are low abundance. Overall, our new pY enrichment reagent simplifies and improves the accessibility of pY-enrichment methods, which will enable more in-depth studies of pY signaling in cell culture and tissue samples for human health and broader biology.

## Results and Discussion

### HaloLink^TM^ bead chemistry simplifies sSrc bead preparation and outperforms NHS chemistry

To improve on our previous pY automated enrichment protocol and develop an easy, cost-effective, and scalable approach for enriching phosphotyrosine, we first condensed the purification and conjugation of sSrc into a single step by fusing a HaloTag N-terminal to sSrc (Figure 1A and 1B and Expanded View Content). Starting from *E. coli* cell pellets, the previous NHS-sSrc bead preparation required 3 days with significant hands-on time (Figure 1C). In comparison, the new Halo-sSrc bead preparation takes only 2 hours and the entire process from bead preparation to enriched pY peptides ready for MS measurement can be completed in the same day. Not only are the Halo-sSrc beads easier to prepare, but they also outperform the NHS-sSrc beads. When benchmarked side by side on a tryptic digest of pervanadate-treated HeLa cell lysate, Halo-sSrc beads yielded more pY peptide identifications and higher signal than NHS-sSrc beads (Figure 1D and EV1). Further, the Halo-sSrc beads maintain higher sensitivity than NHS-sSrc over a range of peptide input amounts (Figure EV2). We attribute the superior performance of the Halo-sSrc beads to multiple factors. The sSrc domain is likely stabilized by the Halo fusion protein^73^ and less prone to destabilization during the quick and gentle conjugation. Further, HaloTag enables single-point conjugation of sSrc to the beads in contrast to NHS beads which react non-specifically to exposed lysines, including a lysine that is part of the pY-peptide recognition interface, which likely impairs peptide binding (Figure EV3).

**Figure 1.**
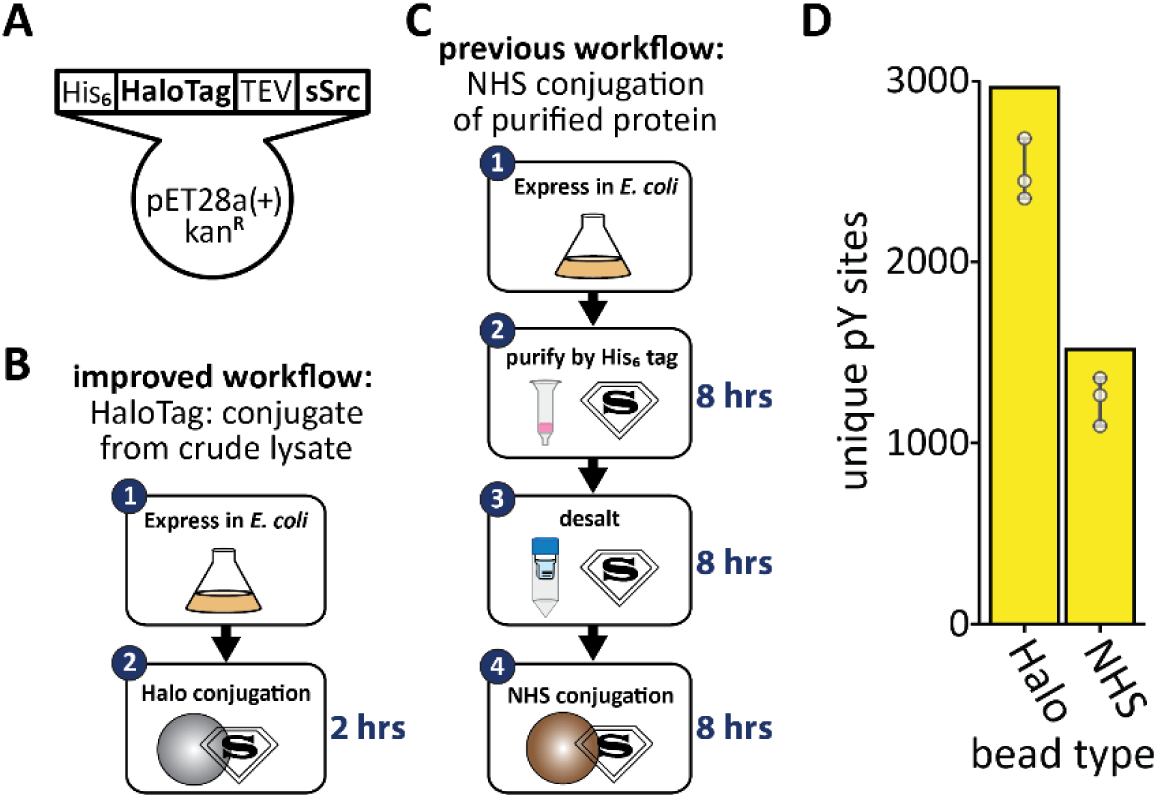
Halo-sSrc is easy to prepare and outperforms NHS conjugation chemistry. **A.** Construct design of Halo-sSrc within the *E. coli* expression vector. From N- to C-terminus, the translated protein under the control of an IPTG inducible promoter consists of a His_6_ tag, serine-serine-glycine linker, a HaloTag (297 amino acids, 33 kDa), serine-glycine-serine-glycine linker, tobacco etch virus (TEV) protease cleavage site (ENLYFQ), serine-glycine-serine linker, and the sSrc protein (109 amino acids, 12.4 kDa). **B.** Schematic of the improved bead preparation protocol where purification and conjugation of Halo-sSrc are performed in a single step by mixing Magne HaloTag beads with crude *E. coli* lysate containing overexpressed Halo-sSrc and washing away unbound material with several PBS washes. **C.** Schematic of the previously published bead preparation workflow requiring pre-purification of His_6_-sSrc by metal affinity chromatography, protein desalting, and spin concentration prior to conjugation to magnetic NHS-activated beads^65^. **D.** Performance of Halo-sSrc beads compared to NHS-sSrc beads for their ability to enrich pY peptides from 1 mg of pervanadate-treated HeLa cell digests in triplicate. Points indicate the number of unique pY sites detected in each replicate and the bars indicate net numbers of unique pY sites detected across all replicates.

The Halo-sSrc beads offer ten-fold cost savings relative to the commercial pY antibody and five-fold savings relative to the previously described NHS-sSrc beads^65^. This cost reduction is critical for large scale multi-condition pY experiments, as required in systems-level studies. Several factors contribute to the cost-savings of Halo-sSrc beads (Figure EV4). First, reduction of bead preparation time from 3 days to 2 hours cuts labor costs. Second, only half the amount of Halo-sSrc bead slurry is required compared to NHS-sSrc^65^, further cutting cost per sample (Figure EV5). Third, at the time of writing, the Promega Magne® HaloTag beads were significantly cheaper than the Pierce™ or Cube Biotech NHS-activated magnetic beads. Fourth, the Halo-sSrc beads do not require additional reagents such as metal affinity resin for His-tag protein purification or centrifugal filters for buffer exchange and protein concentration. The cost of preparing enrichment beads for 96 samples, considering both reagents and labor, was estimated at $551 for Halo-sSrc beads compared to $2,554 for NHS-sSrc prepared according to our 2023 publication^65^ or $5,172 for Cell Signaling Technology® p-Tyr-1000 MultiMab® magnetic bead conjugate, (Figure EV4). To facilitate implementation of R2HaPpY in other laboratories, we provide a detailed protocol and KingFisher method steps (Expanded View Content).

Taken together, the Halo-sSrc reagent improves accessibility and affordability of pY peptide enrichments by omitting pre-purification of superbinder and improves performance.

### Enrichment with Halo-sSrc is robust to various peptide concentrations

Once we established a simplified pY enrichment reagent preparation strategy, we evaluated the procedure with various peptide concentrations. Most pY enrichment protocols start with a few milligrams of input peptides^40,63,66,69,74^. In addition, high throughput enrichment requires peptide solutions fit in a 96-well plate with capacity for 1 mL of liquid. With this scale in mind, we explored the enrichment performance of Halo-sSrc beads on peptides from pervanadate-treated HeLa cells at concentrations ranging from 0.5 mg/mL to 4 mg/mL (Figure 2). Across all peptide concentrations, enrichment efficiency was over 99% by intensity with more than 3300 pY sites detected per replicate and the measurements were highly reproducible with median CVs less than 25%. The number of pY sites (Figure 2A), reproducibility (Figure 2B), and enrichment efficiency (Figure 2C) were similar across concentrations. We also tested whether peptide concentration had an impact on enrichment success for tryptic digest of non-pervanadate treated HeLa cells, where we expect less tyrosine phosphorylation. We tested 4 mg total input at both 1 mg/mL and 4 mg/mL concentrations in duplicate. We observed slightly more unique pY sites and better replicate reproducibility at higher peptide concentrations (Figure EV6). The enrichment efficiency was 99% for both peptide concentrations. In conclusion, we found the procedure is robust to both dilute and concentrated peptide solutions in both high and low pY-containing samples. This enables flexibility of peptide input amount within the high-throughput 96-well format.

**Figure 2.**
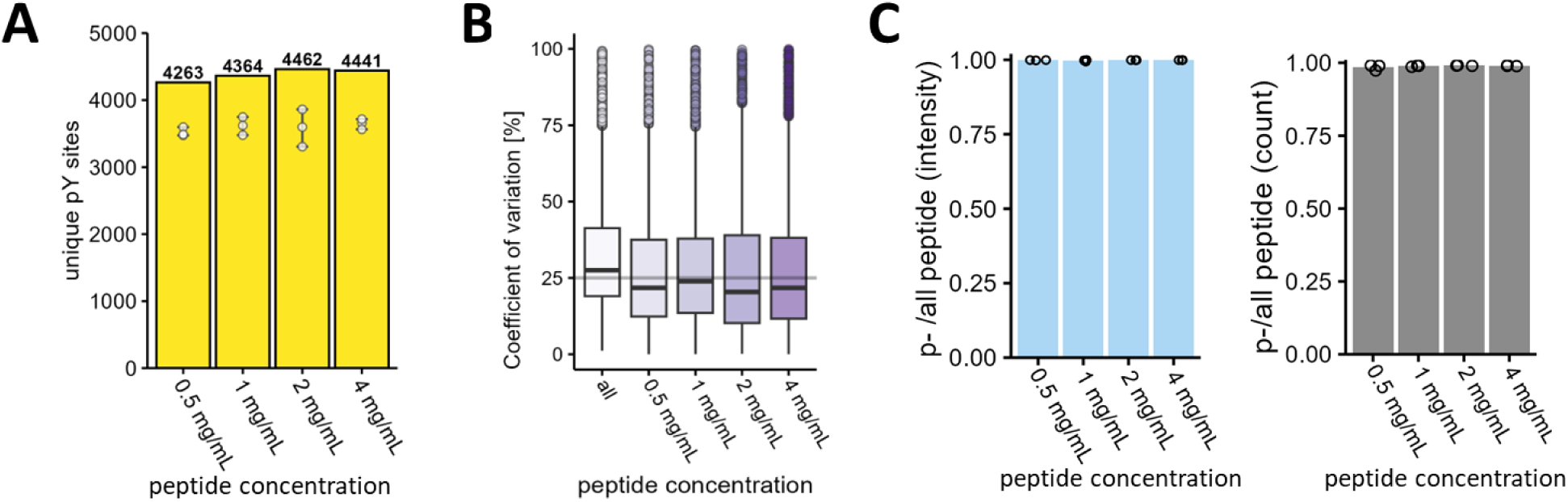
Enrichment with Halo-sSrc is robust to various peptide concentrations. **A.** Bar plots show counts of unique pY sites aggregated from three replicates across four different peptide input concentrations keeping total peptide amount constant at 0.5 mg per replicate. Peptides came from pervanadate treated HeLa cells. Points indicate counts of unique phosphosites per replicate at each of the four peptide input concentrations. Whiskers indicate the range of replicate measurements. Height of bar indicates net unique pY sites from all three replicates. **B.** Boxplots indicate distributions of percent coefficient of variation of pY site intensities between triplicate measurements within a peptide input concentration. A combined percent coefficient of variation across all samples is included at left in white. Intensities of individual phosphosites were summed from all precursors with confident site localization. Intensities were median normalized globally prior to analysis of intensity variation. The horizontal line indicates 25% CV for reference. **C.** Barplots indicate sample purity defined by relative enrichment of phosphopeptides. Ratios indicate the relative intensity (left) or count (right) of phosphopeptides relative to all peptides detected. The height of bars indicates average enrichment efficiency across triplicates and points indicate enrichment efficiencies of individual replicates.

### Halo-sSrc beads are stable for ≥ 4 months

Preparation of large batches of shelf-stable affinity reagents is important for a robust enrichment method that can enable large-scale pY signaling studies. Therefore, we tested the performance of Halo-sSrc beads following four-month storage at 4°C in PBS pH 7.4, alongside freshly prepared Halo-sSrc beads in triplicate. To facilitate quality control throughout method development and reagent characterization, we used a yeast lysate with high levels of tyrosine phosphorylation generated by heterologous expression of an inducible viral Src (vSrc) kinase^75^. We used this sample to test the capacity of different bead batches as well as their shelf life as the sample is scalable and cost-effective. Typically, from only 0.25 mg of yeast lysate input, we enrich over 3000 pY peptides (Figure EV7A and B).

In tests of bead shelf life, the four-month-old beads displayed comparable specificity and capacity as the fresh beads, with all samples enriching an average of 3,400 unique pY peptides per replicate and showing similar intensity distributions (Figure EV8). We also used the yeast lysate with high pY to qualify new batches of beads. Using the vSrc lysate, we tested the quality of 7 independently prepared bead batches. In all batches, we observed a minimum of 3,000 unique pY peptides and enrichment efficiency above 98% (Figure EV7C, D). We conclude that the preparation of Halo-sSrc is reproducible and that Halo-sSrc retains pY peptide binding capacity when stored at 4°C for at least 4 months.

In addition to allowing quality control on bead production, the control peptides can also be used during high throughput experiments to ensure high enrichment efficiency, optimize LC gradients and mass spectrometer methods, as well as monitor LC-MS performance across measurements. The sample is significantly cheaper and easier to produce compared to positive control samples generated from pervanadate treated cancer cells used in other studies^15,30,33,34,38,40,56,60–66,76^.

### R2HaPpy deeply measures phosphotyrosine signaling response in EGF stimulated HeLa cells

Next, we applied our R2HaPpY protocol to investigate epidermal growth factor receptor (EGFR) signaling. Dysregulation of tyrosine kinase signaling networks drive disease, especially cancer, and have become prominent drug targets^10–12,43,44,77–80^. EGFR signaling networks are rewired in numerous cancers, particularly non-small cell lung cancer, and both targeted and combination therapies that target the EGFR pathway are still being explored^10,19,20,80–83^. EGFR signaling has been previously studied using phosphoproteomics^43,44,47,84,85^. But given its importance in cancer and the potential value of studies of drug mechanism of action focused on the EGFR pathway and/or related phosphotyrosine signaling^10–13,16,17,19,80,86–92^, we sought to demonstrate the sensitivity of our method on this well-studied pathway to demonstrate we capture known signaling events and to uncover potentially novel regulated pY sites. HeLa cells were stimulated with EGF and samples were harvested before stimulation and at 1 min, 3 min, 5 min, and 15 min after EGF addition (n = 6 biological replicates). In total, we identified 1,651 unique pY sites with anywhere between 763 and 1189 unique pY sites per condition (Figure 3A, Dataset EV1). Untreated cells showed the lowest unique pY sites while 1 min EGF treatment showed the most. Measured unique pY sites remained above 1,100 through 15 min EGF treatment. Detection of close to 1000 pY sites per replicate from only 1.3 mg of sample material using label-free data dependent acquisition represents a significant improvement compared to published pY enrichment methods which often detect lower numbers of unique pY sites per replicate from higher sample input (147 – 720 pY sites per replicate, from 2 mg to 5 mg per replicate)^16,17,64,74^. This demonstrates the method is capable of profiling the pY proteome at great depth from low sample input.

**Figure 3.**
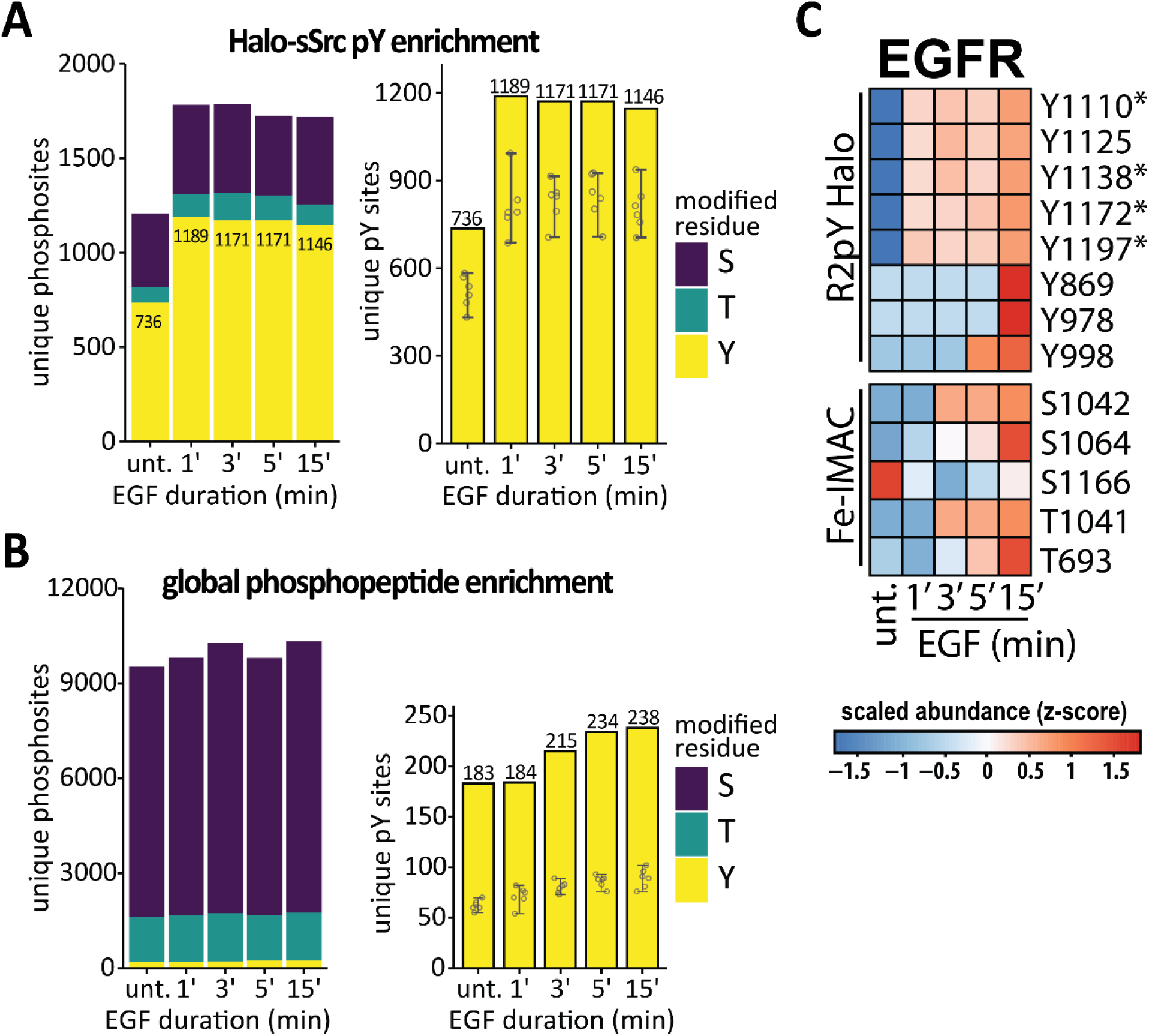
Overview of EGF stimulated HeLa phosphopeptide enrichments. **A.** Stacked bar plot indicates counts of unique phosphosites following pY specific enrichment using Halo-sSrc reagent. Identity of phosphorylated amino acid is indicated by purple (pS), teal (pT), or yellow (pY). Numbers indicate counts of unique pY sites across all six replicates. Right plot only shows counts of unique pY sites. Points indicate unique sites per replicate, while whiskers indicate range of individual replicates. The height of the yellow bar indicates the total unique pY sites across all six replicates. **B.** Stacked bar plot indicates counts of unique phosphosites following global phosphopeptide enrichment with Fe^3+^ metal affinity (Fe-IMAC). Identity of phosphorylated amino acid is indicated by purple (pS), teal (pT), or yellow (pY). Right plot shows counts of only unique pY sites. Numbers indicate counts of unique pY sites across all six replicates. Points, whiskers, and height of yellow bar indicate same as in panel A. **C.** Heatmap representing temporal response of EGF-regulated phosphosites on epidermal growth factor receptor (EGFR). Phosphosite intensities were averaged across replicates and scaled by z-score. EGF-regulated pY sites from R2HaPpY are displayed on top while pSer and pThr sites from global phosphopeptide (IMAC) enrichment are displayed on bottom. Asterisks indicate the four pY sites on EGFR detected in both Fe-IMAC and pY enrichment.

We also enriched the global phosphoproteome using Fe^3+^ immobilized metal affinity chromatography (Fe-IMAC) and found 80% less unique pY sites per condition compared to the Halo-sSrc pY enrichment (Figure 3B). This translates to limited pY site coverage across many proteins. For example, global phosphopeptide enrichment only quantified 4 regulated pY sites on EGFR while Halo-sSrc pY-specific enrichment quantified 8 regulated pY sites on EGFR (Figure 3C). Besides the significant increase in unique pY sites, we found the pY-specific enrichment with Halo-sSrc better distinguished EGF treatment durations compared to the global phosphopeptide enrichment (Figure EV9). Overall, the R2HaPpY method captured a time-resolved EGF signaling response at greater depth than previous studies, in part due to a better and cheaper enrichment reagent which enabled analysis of more replicates.

To analyze the response of pY sites to EGF treatment, we required pY sites to be measured in at least 50% of replicates of at least one condition. After filtering the data, 1,122 unique pY sites remained, with a range of 629 – 1001 unique pY sites reproducibly measured per condition. All subsequent analyses refer to this filtered dataset.

To identify EGF responsive pY sites in our dataset, we performed moderated t-tests on pY site intensities between untreated and EGF-treated cells at each time-point. To account for sites missing from all replicates of a condition, we imputed missing intensities as described in Methods. As expected for short treatment durations, the proteome minimally changed and thus phosphosite abundances were not adjusted (Expanded View Content). Relative to untreated, 78% of well measured pY sites showed a statistically significant intensity change in at least one time-point (moderated t-test, Benjamini-Hochberg adjusted *q* ≤ 0.05). At any individual time-point, over half of the measured pY sites were differentially regulated compared to the untreated samples. Most pY sites were upregulated (48-52%), while only a few were downregulated (7-9%), consistent with previous observations^13,14,17,39,64,93,94^. Importantly, identification of both changing and unchanging pY sites indicates the R2HaPpY enrichment is capable of discerning perturbation specific signaling response.

### R2HaPpY captures more EGF responsive pY sites than previously detected

Next, we wanted to compare the pY EGF responsive sites we detected to other studies and databases. Recently Jayavelu *et al.* (2024) developed EasyAb^74^, an antibody-based enrichment method which enabled them to quantify more than 1000 pY sites from label free, unfractionated analysis of EGF stimulated HeLa cells. Compared to this method, R2HaPpY quantifies more pY sites per replicate and identifies ten times more EGF regulated pY sites (878 regulated sites using R2HaPpY compared to 84 regulated sites with EasyAb). We observe many of the same regulated sites that Jayavelu *et al.* (2024) detected, including known pY sites on EGFR (Figure 3C) and on Met, Ptpn11, Gsk3b and Stat5 (Figure EV10A). This also includes a novel pY site they identified with the EasyAb method (Figure EV10B). Importantly, we identified these regulated sites and many more using approximately half the input material and at 20-times less reagent cost.

We also compared our data to a time course study of early EGF signaling in MCF-10A cells by Reddy *et al.*^13^ that used a combination of three anti-pY antibodies and TMT multiplexing for quantification. Reddy *et al*. quantified 159 pY sites using a 100 nM EGF stimulation over an 80 second time course. While we quantified more than ten times the number of pY sites, we did observe many of the sites detected by Reddy *et al.* with over 50% of the pY sites they detected also being quantified in our dataset (91/159 pY sites).

To more broadly benchmark how well R2HaPpY captured pY signaling events in response to EGF stimulation, we compared the regulated pY sites we identified to previously annotated EGF responsive pY sites and proteins in the PhosphoSitePlus, PTMSigDB, Cell Signaling Technologies (CST), and Wikipathways EGFR signaling (WP437) databases. First, we analyzed the database overlaps at the pY site and protein level (Figure EV11A and B). Surprisingly, there was low agreement between databases at both protein and pY site levels. Only 18% of pY sites were shared between PhosphoSitePlus and PTMSigDB, while only 4% of proteins with annotated roles in EGF signaling were shared between all four databases. Over 75% of proteins were unique to one database. The poor overlap between databases is likely due to different curation strategies. For example, PhosphoSitePlus includes low- and high-throughput studies from many different cell and tissue types and for a site to be annotated as EGF responsive, the phosphosite only needs to be observed as regulated once. In contrast, PTMSigDB contains EGF responsive sites and proteins that are curated from published datasets and requires a consistent direction of change reported by more than one study, in order for a site to be annotated as EGF responsive.

To create a broad and comprehensive signature for EGF stimulation, we combined the data from all four databases encompassing 602 regulated pY sites on 439 proteins (Figure EV11A and B). Of the annotated EGF regulated pY sites, we find close to 30% similarly regulated in our dataset (Figure 4A). Additionally, close to 50% of the regulated pY sites we identified were on proteins that have been annotated as part of the EGF response pathway. Figure EV11C shows the temporal responses of sites that align with known, annotated EGF-responsive pY sites and proteins. The overlap between EGF-regulated pY sites identified in our dataset and previously identified pY sites demonstrates that R2HaPpY can effectively capture known pY signaling.

**Figure 4.**
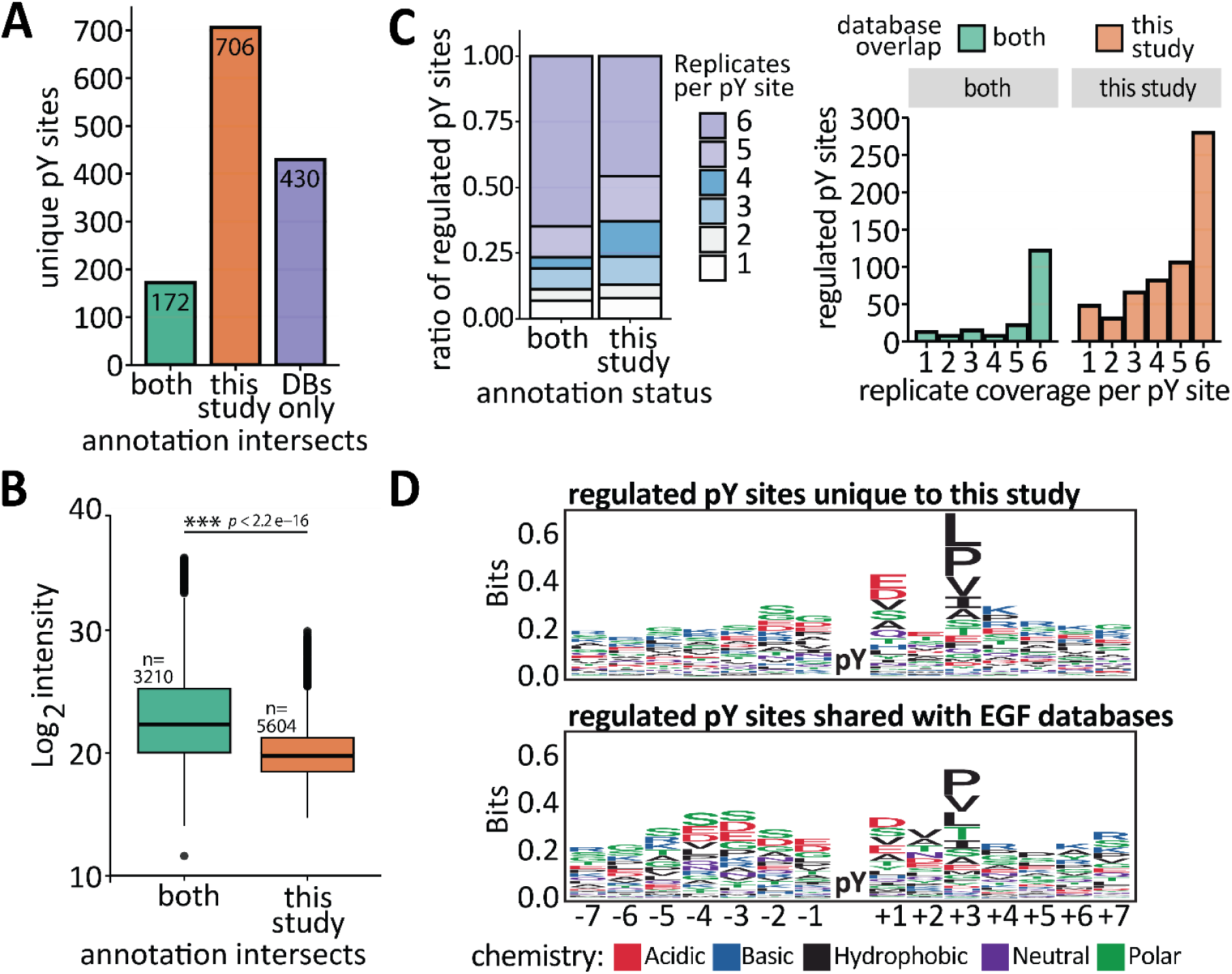
Examination of expanded phosphotyrosine profiling by R2HaPpY. **A.** Barplots indicate counts of unique pY sites measured in this study that were previously annotated by at least one site-level database (green) or not previously annotated by either database (orange). For comparison the purple bar indicates pY sites annotated in either database but not observed significantly regulated in this study (purple). **B.** Boxplots indicate distributions of pY site intensities for sites previously annotated in EGF signatures from either PSP^71^ or PTMSigDB^104^ (green) relative to pY sites lacking database annotation (orange). Only sites measured in all 6 replicates of a condition were included to limit the comparison to well measured sites. This equates to comparison between 132 previously annotated pY sites and 281 pY sites unique to this study which show EGF-dependent regulation. We treated replicate measurements separately, leading to comparison of 3,308 observations relative to 5,535 observations, respectively. Welch’s two sample t-test indicated that average intensity of pY sites unique this study was 4.5- to 5.6-fold lower abundance than sites with prior database annotations within a 95% confidence interval (Welch’s t-test, *p* < 2.2e^-16^). **C.** Left: Distribution of replicate coverage for regulated pY sites, separated by their membership in databased EGF response signatures. Phosphotyrosine sites observed in five or all six out of six replicates of a condition are colored purple. Replicates observed in three or four out of six replicates are highlighted in blue. These pY sites are highlighted to indicate relative proportions of sites that are at higher risk of being missed in a three-replicate experiment. Lastly, pY sites observed in only 1 or 2 replicates of a condition are colored white and gray for reference. Right: Bar plot showing the distribution of replicate coverage across regulated pY separated by their inclusion in databased EGF signatures. X-axis indicates number of observed replicates while the y-axis indicates number of unique EGF-regulated pY sites at each observation extent. **D.** Top: Sequence logo of EGF regulated pY sites identified in this study that were not present in site-level databases (PhosphoSitePlus or PTMSigDB). Logos were made in R using the ggseqlogo package. Height of each letter indicates relative amino acid conservation at each position relative to other peptides in this set. Total number of sequences was 706. Bottom: Sequence logo of EGF regulated pY sites identified in this study that were also present in site-level databases (PhosphoSitePlus, PTMSigDB). Total number of sequences was 170.

### R2HaPpY captures novel, low abundance EGF-regulated pY sites

In addition to capturing known EGF-responsive sites, we also identified many novel sites. Compared to PhosphoSitePlus and PTMSigDB, 80% of the EGF-regulated pY sites that we identified were not previously annotated (Figure 4A). We hypothesized that R2HaPpY captured more EGF-regulated pY compared to previous studies for three main reasons. First, R2HaPpY captures lower intensity sites that are absent in previous experiments. Comparing the intensity of novel regulated pY sites identified here to the sites previously measured in either PhosphoSitePlus or PTMSigDB, we find previously observed sites are >4-fold more abundant than the novel sites we detect (Figure 4B).

Second, R2HaPpY allows us to identify many novel sites because the cost effectiveness and scalability of our approach allows us to prepare and measure more replicates than was possible in previous studies. When we compared the number of replicate observations for novel and previously annotated pY sites, we found that novel EGF-regulated pY sites were more often seen in only three or four of six replicates (Figure 4C). Additionally, 65% of previously annotated pY sites were detected in all replicates, compared to only 46% of novel pY sites.

Third, the sSrc pY superbinder used in R2HaPpY has a unique sequence preference compared to the pY antibody used in the majority of previous studies, which allows us to enrich new, previously unannotated sites. Sequence logos generated from novel regulated pY sites identified only in this study demonstrate the pY superbinder has sequence specificity for glutamic acid or aspartic acid at +1 and leucine, isoleucine, proline, or valine at +3 (Figure 4D), which is consistent with our previous results using the pY superbinder^65^. These results suggest that the pY superbinder sequence preference, combined with our method’s ability to enrich lower intensity pY sites from more replicate samples, allows us to detect novel EGF-responsive pY sites that were previously unannotated.

### R2HaPpY captures previously unseen pY sites to facilitate EGF signaling hypothesis generation

Out of the novel EGF-regulated sites detected with R2HaPpY, 70 pY sites have not previously been reported at all in PhosphoSitePlus, which already currently contains 39,118 pY sites (Dataset EV2). To better understand why these sites have never been observed, we examined the peptides and found seven novel pY sites occurred on peptides with N-terminal clipping and/or acetylation. These included LIMD1-pY4, TKT-pY4, NECAP-pY6, RAB14-pY6, STX12-pY9, PLCG2-pY13, and GRB10-pY15. It is possible these sites were missed in earlier studies due to database search parameters.

To investigate the biological relevance of these 70 novel pY sites, we mapped the sites onto their respective proteins and investigated them in the context of the four different EGF signaling databases described above. We found that 19 of the novel sites were on proteins previously implicated in EGF signaling response (Figure EV11C). One novel pY site at residue Y15 on GRB10, an important scaffold protein involved in growth and proliferation^95–101^, is the first evidence of a post-translational modification within the first 67 residues and may affect tetramerization and downstream signaling^96^. Another novel EGF-regulated site, Y4 on transketolase protein TKT, could modulate nuclear localization. While TKT is an enzyme in the pentose-phosphate pathway, its nuclear localization and interaction with EGFR, MAPK3, and nuclear import machinery requires Y4, even though no PTM has previously been identified on these residues^102^. Importantly, nuclear localization of TKT has been associated with high metastatic potential in hepatocellular carcinoma and therefore modulating this PTM site could present a new therapeutic avenue.

We also identified previously unseen pY sites on proteins not previously annotated in EGF response databases. For example, we identified a number of novel pY sites on proteins known to modify other proteins or participate in PTM crosstalk^6,103^, such as the first documented pY site on the protein modifier SUMO1 (pY21), the second documented pY site on STAMBPL1 (pY238), a zinc metalloprotease that specifically cleaves Lys-63-linked polyubiquitin chains, and the second documented pY site on the E3 ubiquitin ligase RAD18 (pY117). The involvement of these novel sites within EGF signaling is currently speculative but demonstrate how a deep systems level analysis of pY signaling using our method can uncover new, potentially therapeutically relevant pY sites and can drive hypothesis generation for further experiments.

### Uncovering the phosphotyrosine temporal response to EGF

After mapping known and novel pY signaling events that occur in response to EGF stimulation, we wanted to examine the temporal response of EGF-regulated pY sites. To do this, we performed fuzzy c-means clustering of individual regulated pY sites across time points and uncovered four unique clusters that describe the temporal EGF-signaling pY response (Figure 5A), which are summarized in Figure 5B. Regulated pY sites fell into four clusters: 1) sites that rapidly increase at 1 minute EGF stimulation, and stay high through the full time course; 2) sites that increase after 1-3 minutes of EGF stimulation followed by gradual decay after 15 min; 3) sites that steadily increase across the entire time course; and 4) sites that steadily decrease within the first 5 minutes of EGF treatment (Figure 5B). We show the intensity traces of all pY sites colored by their correlation with the cluster trend in Figure EV12A. The majority of previously observed pY sites fall into cluster 1 (60%; 103 of 172) (Figure EV12B), which is likely because pY sites in cluster 1 have higher overall intensities (Figure EV12C). One representative pY site that has been previously observed and one novel site from each cluster are shown in Figure 5C. As expected, novel sites in each cluster are lower intensity compared to previously observed pY sites.

**Figure 5.**
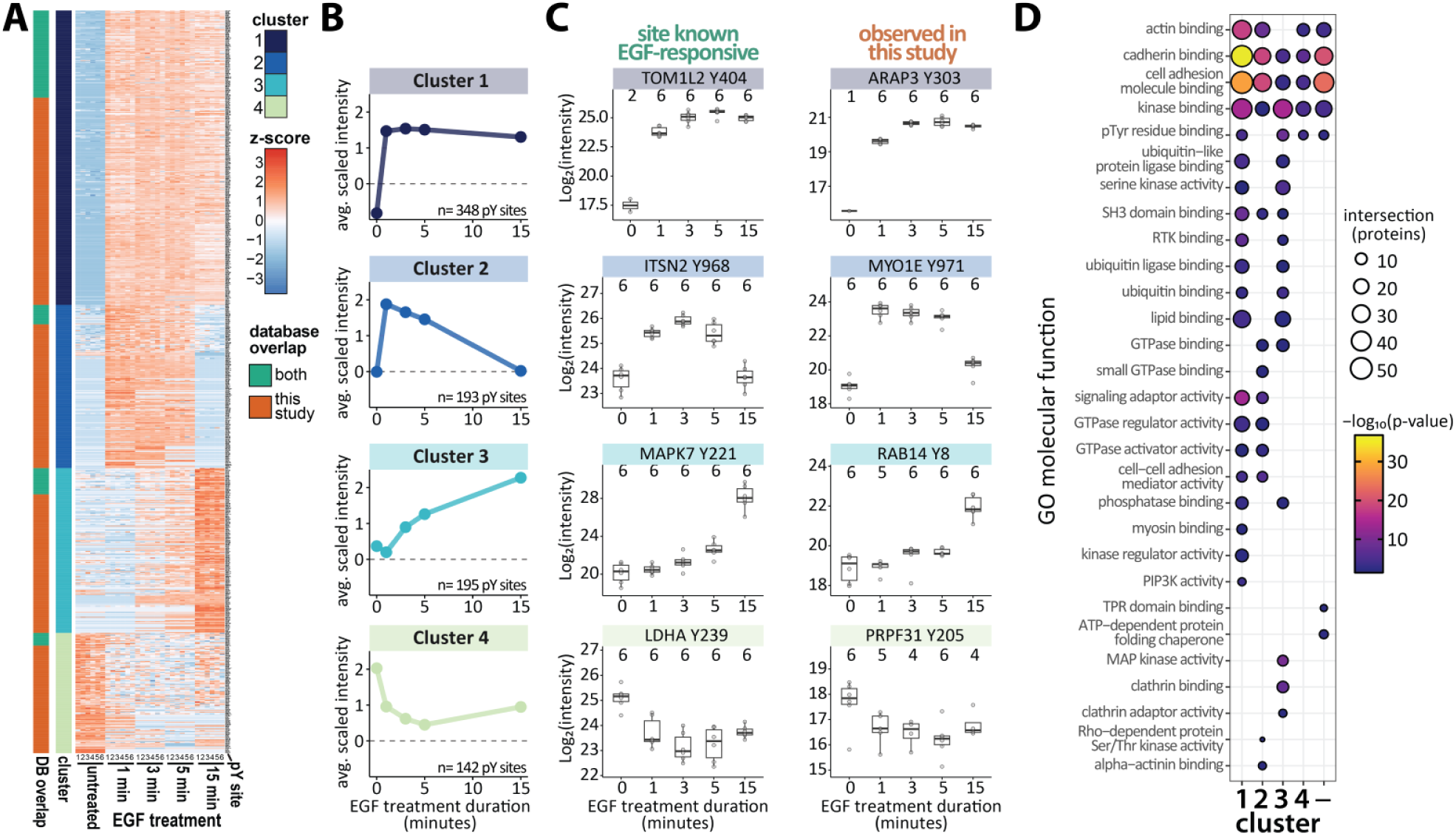
Temporal response of EGF-regulated pY sites and protein functional enrichments. **A.** pY site responses fall into 4 distinct clusters. Clusters were assigned by fuzzy c-means soft clustering method. Only pY sites with a significant abundance change relative to untreated were included in the heatmap. Heatmap shows scaled intensities of individual pY sites (rows) per biological replicate and EGF treatment duration (columns). Rows are organized by cluster and then by overlap with prior database annotation. Rows are annotated to indicate overlap with prior database annotation and cluster membership (left). Database overlap annotation is green if the pY site has been annotated in either PhosphoSitePlus or PTMSigDB EGF response signatures and orange if the site is not in either database. Cluster coloring proceeds from dark blue to light green for clusters 1 through 4. **B.** Line plot indicates average scaled intensities of all pY sites per cluster across EGF treatment durations. Numbers of pY sites belonging to each cluster are listed at the bottom of each plot. **C.** Examples of individual pY sites showing temporal responses to EGF treatment for each cluster. The left column shows pY sites previously annotated as EGF responsive by PhosphositePlus or PTMSigDB. The right column shows pY sites observed in this study without prior annotation in either database. Points indicate individual replicate measurements and boxplots summarize the distribution of measurements per treatment duration. The number of measured replicates is indicated at the top of the plots. **D.** Gene ontology molecular function enrichment results of proteins containing pY sites within each cluster or otherwise not regulated (x-axis dash). Enrichment scores were determined with the gprofiler2^108,109^ tool. Significant molecular functions are listed on the y-axis and cluster is indicated on the x-axis. The size of points indicates the number of proteins within each cluster that match the molecular function term. The color of points indicates the significance of the functional enrichment, where yellow is more significant while blue is less significant. Absence of a point indicates no significant enrichment.

To understand the functions of proteins within each EGF response cluster, we performed gene ontology enrichment analysis, looking at cell component, biological process, and molecular function (Figure 5D and Figure EV13A-B). As expected, proteins involved in cell structure modulation were enriched among all clusters but were most significant in clusters 1 and 2 which reached maximal pY site intensity at one minute, consistent with cell morphology changes in early EGF signaling^13,105^. Additionally, proteins involved in GTPase signaling were enriched in clusters 1 and 2, consistent with the transient proximity of GTPase activators to GTPases during the internalization of the EGFR-Grb2-SOS complex one to five minutes after EGF stimulation^106,107^ (Figure 5D and Figure EV13B,C). We also observed that regulated pY sites on serine-threonine kinases are enriched in cluster 1 and 3 response types (Figure 5D and Figure EV13C), suggesting that pY activation of these kinases either reaches maximal signaling within 1 minute and is sustained or gradually increases through 15 min. Conversely, regulated pY sites on MAPKs were primarily enriched in cluster 3, with gradual increase across the time course.

In the enrichment analyses, we also included pY sites which were not regulated in response to EGF and stayed the same throughout the time course. These non-EGF regulated pY sites were enriched in multiple molecular functions, biological processes, and cellular components, however these sites were uniquely enriched for nucleo-cytoplasmic transport (Figure 5D and Figure EV13A-B). As translocation of proteins is a key part of signaling pathways and often facilitated by pY signaling, this enrichment suggests R2HaPpY detects a broad range of pY sites that could potentially be regulated at longer time points after EGF stimulation, or which could be regulated in response to alternative stimulations. Overall, temporal monitoring of pY sites at a systems level captured lifetimes of signaling responses across a broad range of protein functional classes. These observations underscore the benefits of systems-level view of pY signaling using a sensitive enrichment method.

### Temporal regulation patterns of EGF-responsive pY sites do not correlate with signaling pathway depth

Reconstructing signaling networks is an active area of research which has begun to uncover functional connections that are not represented in linear signaling maps^3,21,110,111^. Importantly, network reconstruction requires measurement of the signaling system under multiple perturbations and time points, which is only feasible with high-throughput methods like ours^2,21^. We therefore examined if our deep and temporally resolved phosphotyrosine dataset can help understand network complexity. To assess this, we investigated whether the four distinct temporal response profiles we observed across the EGF regulated pY sites correlated with the position of each protein within an EGF signaling pathway map, such as WikiPathway 437 (WP437)^112^. If WP437 fully represented the network, we would expect proteins close to EGFR to have rapid response profiles and occur in clusters 1 and 2, while proteins further down the EGF pathway would have regulated pY sites that increase in abundance more slowly, such as the sites in clusters 3. We devised a node depth analysis where proteins in the human EGF/EGFR signaling pathway (WP437) were assigned a depth by tallying nodes while tracing edges extending from EGFR (Dataset EV3).

Regulated pY sites were then ordered by the protein-level node depth and displayed alongside a heat map denoting cluster assignment and scaled pY site intensity at each EGF treatment duration (Figure EV14A). As expected, protein node depth was not clearly correlated with pY site temporal response profiles (Welch’s t-test, *p* < 0.05) (Figure EV14A and B).

This lack of correlation between protein node depth and pY site temporal response contrasts with the common visualization of signal transduction as sequential phosphorylation emanating from the cell surface and instead agrees with other systems-level protein interaction and signaling studies that have suggested signaling networks are not exactly linear^2,3,13,110,111^. Still, only one systems level pY study has pointed to this issue underscoring the need for cost effective and high throughput pY enrichment methods. For example, in response to EFG stimulation, Reddy *et al.* observed rapid tyrosine phosphorylation of cell morphology proteins tensin, cortactin, and plankophilin despite no known direct interaction with EGFR^13^. We observe a similar response of the same pY sites after 1 minute of EGF stimulation, in addition to other pY sites that are often lower abundance on the same proteins.

We highlight several additional examples from our dataset that provide further evidence of signaling network complexity and the challenges in reconstructing it. For example, we examined proteins which we expect to be phosphorylated later in the EGF signaling response based on their node depth, such as the MAP kinases (MAPKs), which mediate diverse cellular functions such as cell growth, adhesion, survival, and differentiation through regulation of transcription, translation, and cytoskeletal rearrangements. We detected 15 EGF-regulated pY sites across 11 MAPKs and two MAPKAPKs (Figure EV15). Nine of these 15 pY sites were at known activation loop positions which showed expected gradual intensity increase without plateau by 15 minutes. Unexpectedly however, two pY sites on targets of the MAPKs, MAPK activated protein kinases MAPKAPK2 pY228 and MAPKAPK3 pY207, rapidly increase in abundance after 1 minute of EGF stimulation, followed by dephosphorylation through 15 minutes. Both of these pY sites, MAPKAPK2 pY228 and MAPKAPK3 pY207, are positioned between known kinase activating pS and pT sites^113^. MAPKAPK2 pY228 is novel and MAPKAPK3 pY207 has no annotated function. Our temporal results suggest a potential role in early response to EGF stimulation and kinase activation, which will be interesting to investigate further.

Similarly, the scaffold protein IQGAP1 is known to bind B-Raf, MEK, and ERK^114^ and had a node depth of 5 towards the bottom of the EGF/EGFR signaling pathway^112^. We hypothesized that regulation of pY sites on IQGAP1 would occur after 1 minute of EGF stimulation, consistent with MAPK signaling timescales. Instead, we saw rapid regulation of two sites, Y1510 that decreased in abundance initially, and Y17 that increased in abundance after 1 min. Other studies have found that MET phosphorylates IQGAP1 at Y1510 and this site was recently found to bind both full-length Abl1 and Abl2^115,116^. This suggests IQGAP1 is involved early in EGF signaling response given Abl1 has a pathway depth of 1. In support of earlier involvement of IQGAP1, others reported IQGAP1 directly associates with EGFR during EGF stimulation and induces S1443 phosphorylation^117^, consistent with our data. Our sensitive pY enrichment opens the possibility to link early tyrosine phosphorylation to later serine phosphorylation in context of IQGAP1’s direct association with EGFR. To our knowledge, this is the first indication that early phosphorylation at Y17 and later phosphorylation at Y1114 may contribute to IQGAPs regulation in context of EGFR scaffolding.

Additional examples include PTPN11, SOS2, STAT3, GAB1, GAB2, PLCG1, PIK3C2B, PTK2B, IQGAP1, and PRKCD, which were assigned node depths between 2 and 6, and at least one pY site on each protein displays immediate intensity increase within 1 min of EGF treatment. Overall, we found that many proteins depicted at the bottom of the pathway experienced maximal pY site modulation within 1 minute of EGF treatment, encouraging exploration of novel feedback mechanisms and other types of network connectivity.

While there were certain pY sites with regulation inconsistent with the location of the protein in the WP437 network diagram and inadequately explained by displayed interactions, we observed several network branches that displayed response profiles generally consistent with pathway depth. For example, proteins involved in actin polymerization such as, VAV, RAC, CDC42, ROCK1, BCAR1, EPS8, and others showed primarily cluster 4 type responses indicative of immediate dephosphorylation and were concentrated to early pathway depths of 1 – 3 (Figure EV14C).

Similarly, early transcriptional modulation mediated by STATs, PIAS3, E2F1 and PCNA showed cluster 1 and 2 response profiles of maximal pY intensity by 1 min followed either by sustained activation or gradual decay, respectively (Figure EV14C). Finally, the later transcriptional modulation mediated by MAP kinases showed primarily cluster 3 response profile consistent with their deeper position in the WP437 network. This suggests that current signaling maps are valuable references to explain a portion of interactions, however with sensitive pY profiling, additional network dependencies and connectedness can be revealed.

### Differential regulation of EGF-responsive pY sites on the same protein

As we were examining temporal pY profiles in the context of the EGF/EGFR signaling network, we noticed that many EGF-responsive pY sites that fall into different clusters occurred on the same protein. When we investigated this further, we found 78% of regulated pY sites on proteins annotated in the EGF/EGFR signaling pathway showed a different response profile compared to at least one other pY site on the same protein (Figure EV14A). Olsen *et al.* observed a similar phenomenon in a global phosphoproteome analysis of EGF stimulation where they found 77% of proteins with an EGF-responsive phosphosite also had at least one additional phosphosite that temporally behaved differently^85^.

The differential response profiles of pY sites on the same protein are further evidence of previously undiscovered feedback loops and network connectivity that is more complicated than what is represented in the pathway diagram. As an example, pY sites in different regions of the adaptor proteins CRK and CRKL show different response profiles (Figure 6A-C). Different domains of the adaptor proteins CRK and CRKL are essential for signal propagation during development, in response to various growth factors, including EGF, and in cancerous transformation^118–130^. In CRK, pY251, a site located in the C-terminal SH3 domain known to engage Abl kinase^119^, reached maximal intensity within 1 minute (Figure 6B). In contrast, two other sites observed in the flexible linker between the SH2 and N-terminal SH3 domains, pY108 and pY136, gradually increased intensity without plateauing by 15 minutes (Figure 6B). It is possible these sites participate in a feedback loop to modulate continued EGFR signal propagation, and our data highlights an area for further study. Additionally, CRK pY136 is affected by gefitinib treatment^131^ showing that we capture sites that are modulated by cancer therapeutics.

**Figure 6.**
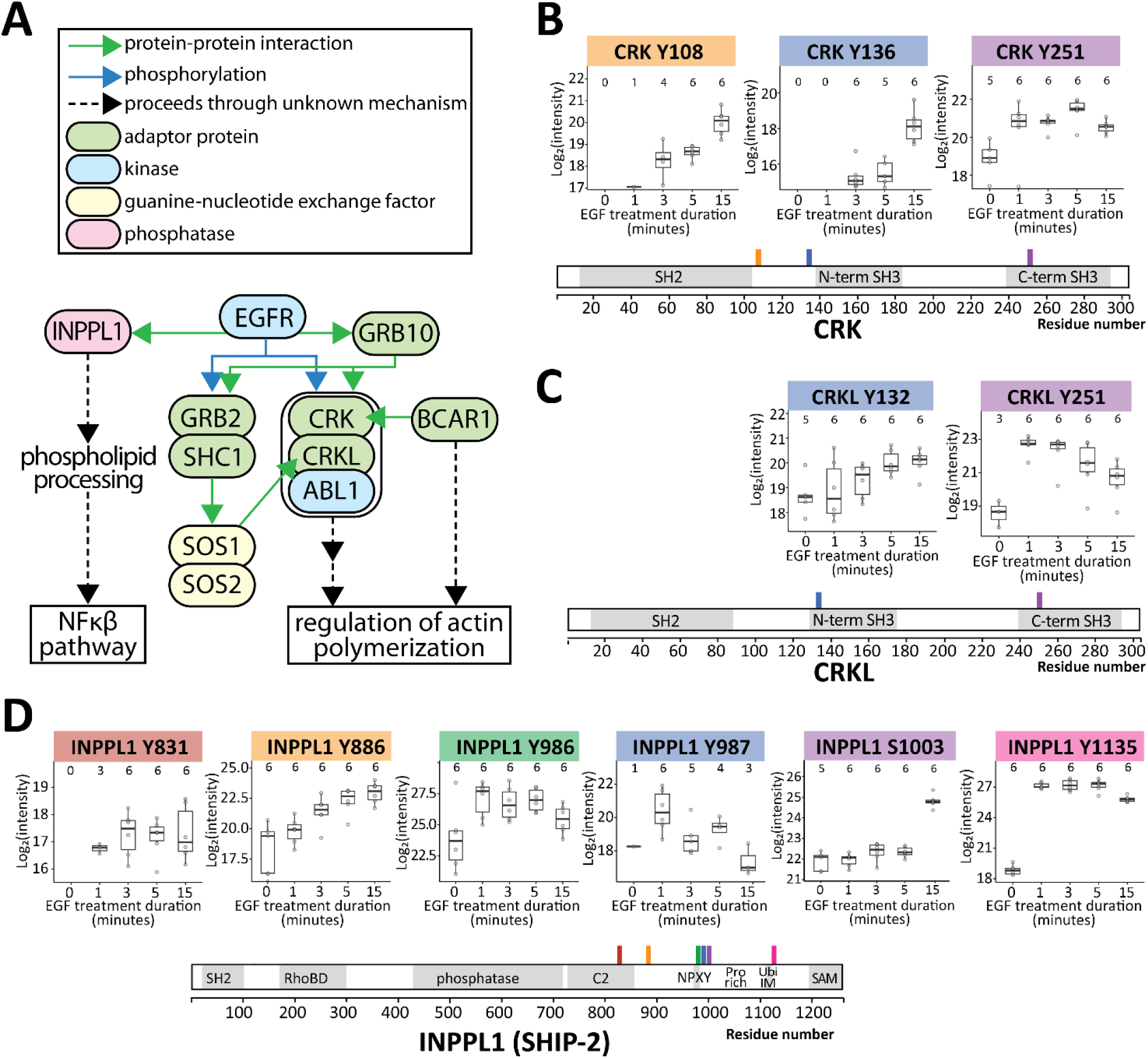
Examples of different pY site temporal profiles within proteins. **A.** Schematic of a subset of EGF/ EGFR signaling adapted from WikiPathways #437. **B.** Boxplots indicate distribution of pY site intensity at each EGF treatment duration for the protein CRK. Points indicate individual measurements per replicate. The total number of replicates for which a pY site was measured per treatment duration is indicated near the top of each plot. The title of each plot indicates the gene name and pY site location in context of the full protein. The background color of the plot title corresponds to the location of the pY site along the schematic of protein domain boundaries indicated. Domain boundaries were adapted from PhosphoSite Plus. **C.** Boxplots indicate distribution of pY site intensity at each EGF treatment duration for the protein CRKL. All other plotting information is the same as for panel B. **D.** Boxplots indicate distribution of pY site intensity at each EGF treatment duration for the protein INPPL1 (SHIP-2). Schematic of domain boundaries was adapted from both PhosphoSite Plus and Le Coq *et al.* (2017, 2021). All other plotting information is the same as for panel B and C. “RhoBD” indicates a putative Rho binding domain. The “phosphatase” domain indicates the 5-inositol phosphatase domain. The “C2” domain indicates the C2 calcium and lipid binding domain. “NPXY” indicates the conserved amino acid motif in which the Y is phosphorylated and is commonly involved in protein trafficking. “Pro-rich" indicates the proline rich region spanning roughly amino acids 930 – 1110. “SAM” indicates the sterile alpha motif.

As another example, CRKL also exhibits regulated pY sites with different temporal profiles. At 1 minute after EGF stimulation, pY251 peaks while pY132 gradually approaches plateau by 15 minutes (Figure 6C). According to PhosphoSite Plus, Y132 is phosphorylated by EGFR, while Y251 is phosphorylated by ZAP70, suggesting we observe convergence of signaling on CRKL across different temporal timescales.

As a third example, the inositol phosphatase SHIP2/INPPL1, which is recruited to EGFR early to downregulate the PI3K pathway^132^, shows divergent pY site regulation indicative of its role as a signaling hub. Specifically, pY986 and pY1135 are directly phosphorylated by EGFR^132,133^ and reach maximal intensity 1 minute after EGF stimulation, consistent with the location of INPPL1 in the EGF/EGFR signaling pathway map (Figure 6A and D). Both EGFR phosphorylated pY sites occur in the proline-rich region of INPPL1, one on the NPXY motif and the other immediately adjacent to the ubiquitin interacting motif^134,135^ (Figure 6D). In contrast, pY886, located closer to the the lipid binding C2 domain, is phosphorylated in response to both Ephrin B1/2 expression^136^ and erlotinib treatment^19^, and shows a slower pY site increase through 15 minutes. Similarly, we observed a much less studied pY site, pY831 located in the lipid binding C2 domain, which also shows a slower pY site increase (Figure 6D). Interestingly, the C2 domain has an activating effect on the lipid phosphatase activity^135^, opening the possibility pY831 may influence this catalytic enhancement.

While further study of specific interactions would be required, our enrichment method can track distinct temporal profiles within individual signaling proteins enabling discernment of pY site modulation under different control mechanisms.

While we focused on proteins annotated in the EGF/EGFR signaling pathway map, we observed a similar trend of multiple distinct temporally regulated pY sites occurring on the same protein across all 878 sites we identified as regulated. Of the 152 proteins with more than one regulated pY site, 66% (100) have pY sites with different temporal response profiles. This finding further supports the notion that post-translational modifications and their regulation should be reported at the site level and not aggregated to the protein level^85,104^. Further, it is exciting that twenty years later we can revisit and validate some of the observations made by Olsen *et al*. at the global phosphoproteome level, but now with significantly increased tyrosine phosphoproteome depth (i.e. 878 versus 53 EGF-regulated pY sites) achieved by a scalable and sensitive pY specific enrichment approach.

### Concluding remarks

R2HaPpY is a method for cost-effective anti-pY peptide bait preparation coupled with automated enrichment in 96-well format on the KingFisher Flex magnetic particle processing robot. R2HaPpY is easy to implement and is sensitive enough to capture pY signaling events at high depth from low sample input. In a simple benchmark of the approach using EGF-treated HeLa cells, we observed both expected and novel EGF-regulated pY sites allowing us to expand what is known about EGFR signaling and enriching current pathway maps. Not only is R2HaPpY sensitive, but the approach is also ten-fold cheaper than commercial antibody-based approaches, and less labor and reagent intensive than other SH2 superbinder approaches, enabling scalability to systems biology studies involving hundreds of samples. We envision this simplified and robust method to elevate investigations of tyrosine phosphoproteome regulation to the scale and regularity seen for global phosphopeptide enrichments, allowing researchers to probe more dimensions of the tyrosine phosphoproteome. Ultimately, improved methods for monitoring site specific tyrosine phosphorylation temporal dynamics will enable deeper understanding of signaling systems.

## Materials and Methods

### Reagent and Tools table

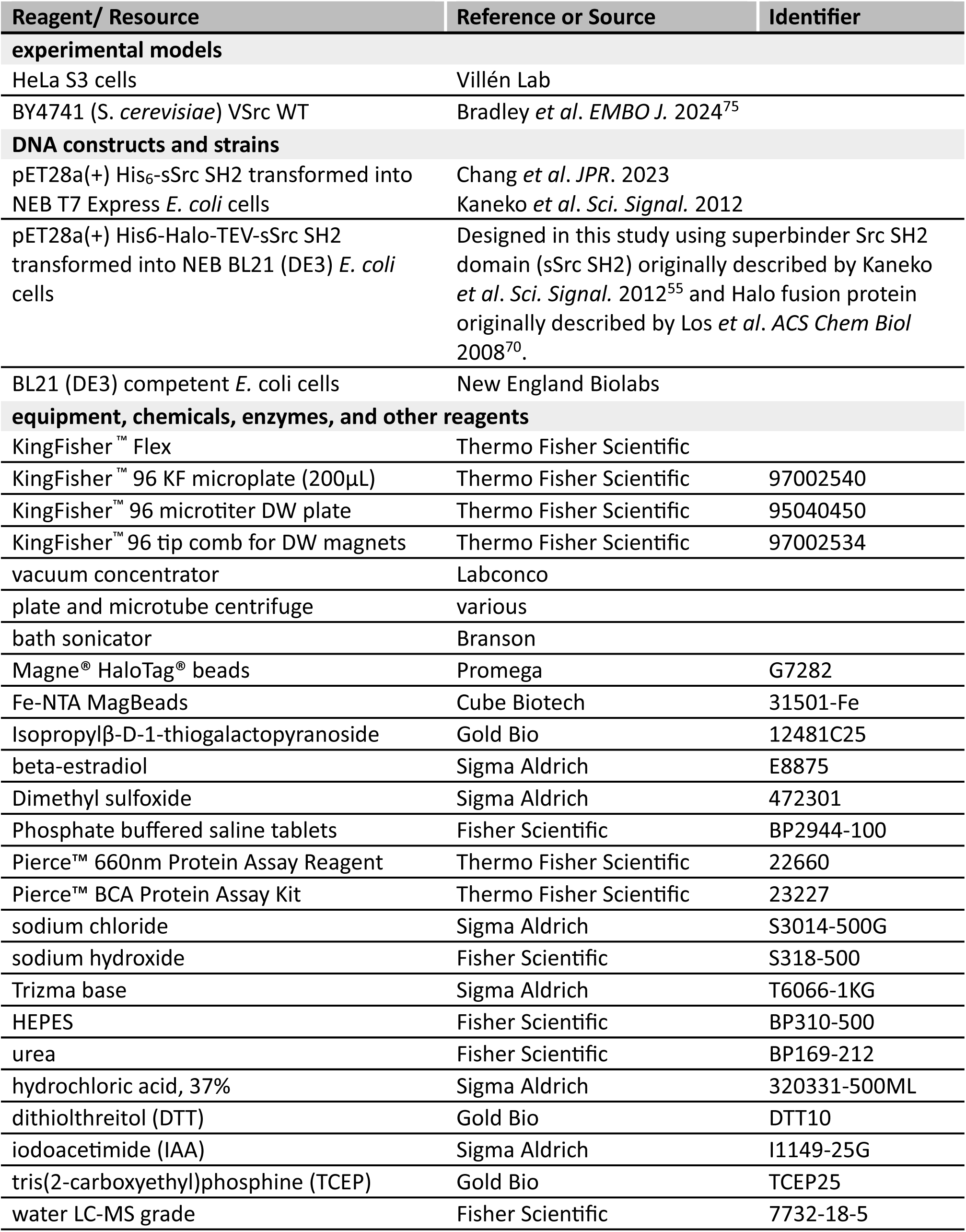
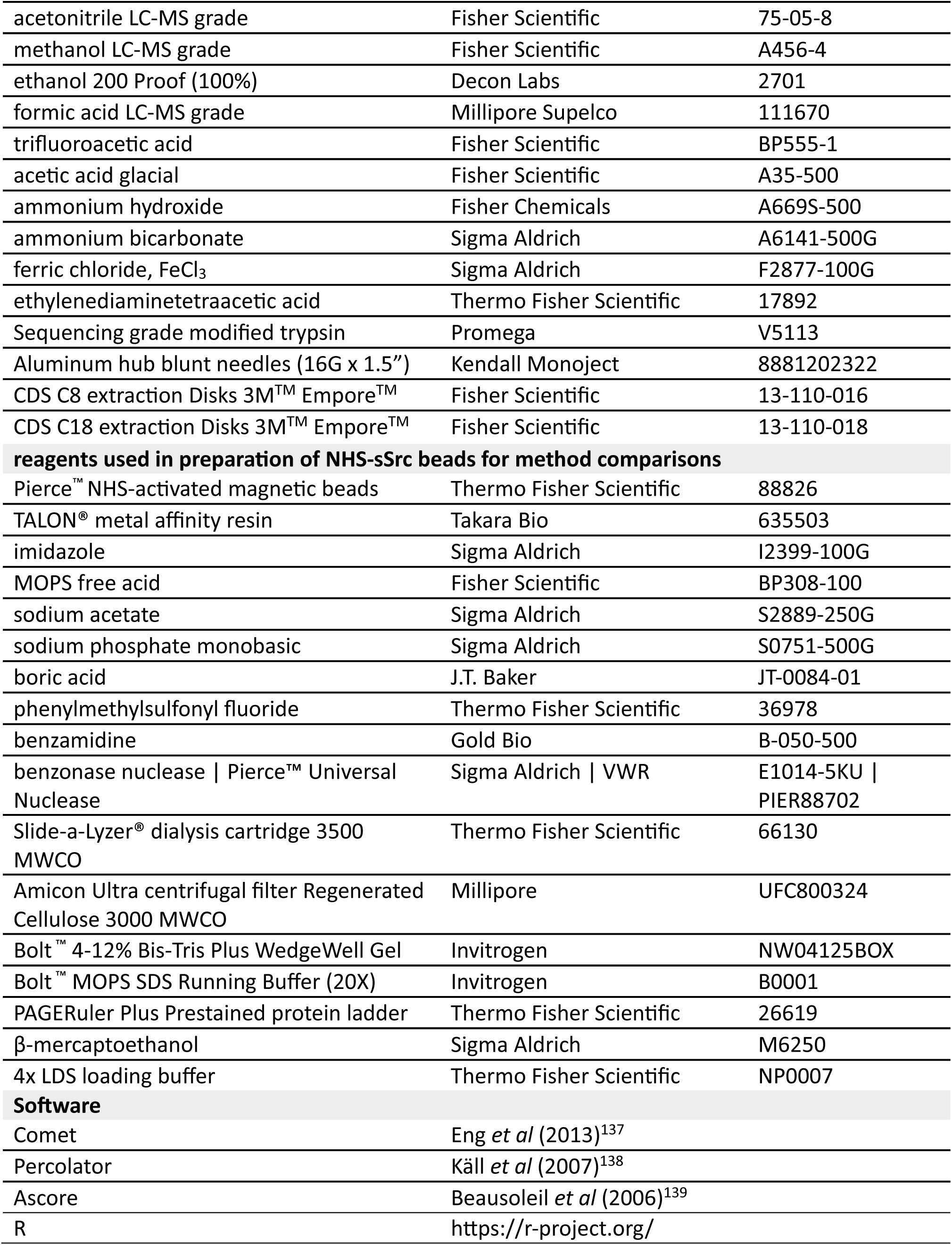

## Methods and Protocols

### Preparation of Halo-sSrc beads Construct design

A HaloTag^TM^ was fused N-terminal to Src SH2 superbinder (sSrc) in the pET28a(+) *Escherichia coli* expression vector. The sSrc protein contains mutations T8V, C10A, and K15L^55^. A Tobacco Etch Virus protease cleavage site was included between the HaloTag and sSrc SH2 domain and a hexahistidine tag was included at the N-terminus. Each component is connected by short serine-glycine linkers. Sequences and plasmid maps for both the full plasmid and open reading frame are shown in Expanded View Content.

### Halo-TEV-sSrc expression

*Escherichia coli* culture (BL21 (DE3) pET28a(+)::His6-Halo-TEV-sSrcV2 Kan^R^) was grown in Luria Broth with kanamycin (50 µg/mL) to OD_600_ of 0.6 – 0.8 then induced with 0.1 mM Isopropyl β-D-1-thiogalactopyranoside (IPTG) for 3 – 4 hours at 37°C with 225 rpm shaking. Cells were pelleted, washed in phosphate buffered saline (PBS), aliquoted into 50 mL culture equivalents, and stored at -80°C.

### Halo-TEV-sSrc lysis and conjugation

Frozen *E. coli* cell pellets containing Halo-TEV-sSrc were resuspended in 1X PBS pH 7.4 supplemented with 0.1 mg/mL lysozyme and 12 – 25 U/mL benzonase nuclease at a ratio of 4 mL buffer to 50 mL culture equivalent of cell pellet. Cell suspension was rocked at 4°C for 5 – 10 min to allow lysozyme and benzonase activity. Cells were sonicated using 11–12-watt output in 6 intervals of 15 – 30 sec with equal rests on ice. Lysate was clarified at 21,000 × *g* for 15 min at 4°C.

Prior to conjugation, Magne® HaloTag® beads (Promega) were equilibrated by washing three times in highly pure (18 Ω) water followed by three washes in 1X PBS pH 7.4. Clarified *E. coli* supernatant was transferred to equilibrated Magne® HaloTag® beads and rocked end-over-end for 45 min to 1 hour at 4°C. Ratio of *E. coli* lysate to magnetic beads was 4 mL clarified lysate (50 mL culture equivalent) per 1 mL of 20% magnetic bead slurry.

Unbound lysate was removed and optionally saved for SDS PAGE analysis as ‘flow-through’. Magnetic beads were washed 6 times in 1 – 2 column volumes (CVs) of 1X PBS pH 7.4. Beads were transferred to a fresh tube after the third wash. Beads were then washed three times in 50 mM Tris, 50 mM NaCl, pH 8.0 prior to pY peptide enrichment or storage at 4°C. Beads can be stored at 4°C for at least 4 months in 1X PBS pH 7.4 or 50 mM Tris, 50 mM NaCl, pH 8.0. To enrich pY peptides, 40 µL of 20% bead slurry was used per sample.

### Conjugation assessment

Successful conjugation of Halo-TEV-sSrc protein to the Magne® HaloTag® beads was confirmed by Tobacco Etch Virus (TEV) protease digest. SDS PAGE analysis of the digest reaction was used to identify presence of the ∼14 kDa sSrc SH2 domain. Briefly, a 50 µL reaction consisting of 10 µL conjugated bead slurry (20% beads ^v^/_v_ in 50 mM Tris, 150 mM NaCl, pH 8.0), 0.2 µg TEV protease (40 µg/mL), 50 mM dithiothreitol, and 10 mM ethylenediaminetetraacetic acid (EDTA) was incubated 16 - 24 hours at room temperature with mixing. Half of the digest (25 µL) was analyzed by reducing and denaturing SDS PAGE using a 4-12% BisTris NuPAGE Bolt gel run at 130 V for 80 min in 1X MOPS/SDS buffer alongside a PAGERuler Plus size marker (Figure EV16). A high intensity 14 kDa band indicated ample sSrc was cleaved away from the HaloTag which remained covalently attached to beads.

### Preparation of NHS-sSrc beads His-sSrc-SH2 expression

The His_6_-sSrc-SH2 construct was prepared as described previously^65^. Briefly, Luria broth containing kanamycin (50 µg/mL) was inoculated with *E. coli* expressing the His-tagged sSrc-SH2 at OD_600_ ∼ 0.1, grown at 37°C with 225 rpm shaking until OD_600_ 0.3 – 0.4 at which point temperature was lowered to 18°C. When cultures reached OD_600_ 0.6 – 0.8 they were induced with 0.1 mM IPTG followed by incubation overnight (18 – 22 hrs) at 18°C with 225 rpm shaking. Cells were harvested by centrifugation at 6000 × *g* for 10 min and 4°C. Cells were washed in 1X PBS pH 7.4 and stored at -80°C.

### His-sSrc-SH2 purification

Frozen cell pellets were resuspended in a lysis buffer containing 1X PBS, 0.1 mg/mL lysozyme, 12 U/mL benzonase nuclease, 1 mM phenlymethylsulfonyl fluoride (PMSF), and 1 mM benzamidine at a ratio of 2.5 mL buffer per 250 mL culture. Cell suspensions were rotated for 10 min at 4°C prior to sonication on ice using four pulses of 20 - 30 sec at 12 watt output with equal rests. Lysate was clarified by centrifugation for 15 min at 21,000 x g and 4°C. Soluble supernatant was further clarified by 0.45 µm syringe filtration using a cellulose acetate membrane. His-tagged sSH2 was immediately purified with TALON® resin (Takara Bio) using a gravity column as described previously^65^. Briefly, purification from 250 mL culture equivalent used 4 mL of 50% TALON ® resin slurry (2 mL dry resin volume) equilibrated in water followed by 1X PBS, pH 7.4. Lysate was mixed with resin and incubated with gentle rotation at 4°C for 60 minutes. Resin was washed with 10 CVs of 1X PBS, pH 7.4, followed by 3 CVs of 1X PBS with 5 mM imidazole then 2 CVs of 1X PBS with 10 mM imidazole. His-sSrc was eluted with 5 CV 1X PBS with 300 mM imidazole. All elution fractions were pooled and buffer exchanged into 50 mM borate, pH 8.5 using 3,000 MWCO centrifugal concentrators at 3000 x g in 10 min increments. Protein was quantified by absorbance at 280 nm using mass extinction coefficient of 0.99.

### NHS conjugation

Pierce NHS-activated magnetic bead slurry (10 mg/mL) was equilibrated to room temperature for 1 hour prior to aliquoting. Beads were equilibrated with 1 mL ice cold 1 mM hydrochloric acid. His-sSrc protein solution at 1.75 – 2 mg/mL in 50 mM borate pH 8.5 was added to beads at 1:1 ratio by volume relative to original bead slurry volume. Beads were mixed gently by inversion for 1 – 2 hours. Beads were collected by magnet and placed into quench/storage solution of 50 mM Tris, 100 mM NaCl, pH 8.0. Beads were washed three times with 50 mM Tris, 100 mM NaCl, pH 8.0 to quench NHS reactivity. The second and third washes were incubated for 30 min and 90 min, respectively. Beads were resuspended to 2 mg/mL in 50 mM Tris, 100 mM NaCl, pH 8.0 for storage at 4°C. Prior to R2pY enrichment, beads were resuspended to 10 mg/mL in 20 mM MOPS-NaOH, 10 mM Na_2_HPO_4_, NaCl, pH 7.2.

### HeLa culture, lysis, and quantification

HeLa S3 cells were cultured at 37 °C and 5% CO_2_ in Dulbecco’s modified Eagle’s medium (DMEM) supplemented with 4.5 g/L glucose, L-glutamine, 10% fetal bovine serum (FBS), and 0.5% streptomycin/penicillin. To generate bulk phosphopeptides for method comparisons, cells were grown to an average of 80% confluency before harvesting. For pervanadate treatment, cells were incubated in serum-free medium for 6 hours prior to addition of 1 mM pervanadate for 15 min, followed by addition of 10% FBS for 15 min. To harvest, cells were rinsed three times quickly with ice-cold PBS then flash-frozen in liquid nitrogen prior to storage at −80 °C. Cells were lysed by scraping in 1 mL denaturing lysis buffer (8M urea, 50 mM HEPES, 75 mM NaCl, pH 8.0) per 15 cm diameter plate. For epidermal growth factor (EGF) stimulation, cells were starved for 20 hours in DMEM containing 0.7% FBS before media was exchanged with DMEM containing 0.7% FBS and 160 ng/mL EGF. Cells were harvested at 1 min, 3 min, 5 min, and 15 min after stimulation. Control plates received no EGF and were directly harvested. Cells were lysed from frozen plates by addition of 8 M urea, 50 mM HEPES, 75 mM NaCl, pH 8.0 Lysate was sonicated twice for 8 sec at 11-12-watt output on ice with 48 sec intermittent rests. Lysate was clarified by centrifugation at 21,100 *x g* and 12°C for 10 min. Soluble lysate was transferred to fresh tubes and quantified by bicinchoninic acid (BCA) assay (Pierce).

### Yeast strains, growth, lysis, and quantification

The yeast strain expressing beta-estradiol inducible v-Src kinase was a gift from the Landry lab^75^. Cells were grown overnight at 30°C with 250 rpm shaking in yeast extract peptone dextrose (YEPD) media. Cells were diluted to OD_600_ of 0.1 in synthetic complete media (1x yeast nitrogen base, 2% glucose, all amino acids) containing 100 nM beta-estradiol prepared in DMSO) and grown to OD_600_ of 0.8 - 1.0. Cells were pelleted, washed with 1x ice-cold PBS, flash frozen in liquid nitrogen, and stored at -80°C. Cell lysis was carried out in 8M urea, 150 mM Tris, pH 8.2 by bead beating four times with 1 min intermittent rests on ice. Lysate was collected then clarified by centrifugation at 4°C and 21,000 × g for 10 min. Protein concentration was quantified by BCA assay.

### Lysate reduction and alkylation

Proteins were reduced with 5 mM dithiothreitol (DTT) at 55°C for 30 min, 1000 rpm mixing. Lysate was cooled to room temperature prior to alkylation with 15 mM iodoacetamide at 24°C and mixing at 700 rpm in the dark. Alkylation was quenched with an additional 5 mM DTT and 700 rpm mixing at 24°C in dark for 30 min.

### Digestion and desalting

Reduced and alkylated protein lysate was diluted five-fold in 50 mM ammonium bicarbonate pH 8.0 containing 1:100 w/w ratio of trypsin (Promega). Proteins were digested for 15 hours at 37°C while shaking at 200 rpm. Digests were acidified to a final pH < 2 with trifluoroacetic acid (TFA). Precipitates were removed by centrifugation at 7000 *x g* for 10 min at 24°C. Peptides were desalted on Waters Sep-Pak 100 mg capacity. Briefly, cartridges were equilibrated with 1 CV methanol, 3 CVs acetonitrile (ACN), 1 CV 70% ACN 0.25% acetic acid (AA), 1 CV 40% ACN, 0.5% AA, and 3 CVs of 0.1% TFA. Peptides were loaded and flow through was reloaded. Cartridges were washed with 3 CVs 0.1% TFA and 1 CV 0.5% AA prior to elution of peptides with 0.8 mL of 40% ACN, 0.25% AA followed by 0.6 mL 70% ACN, 0.5% AA. Eluates were thoroughly mixed, and aliquots made for total proteome measurement, global phosphopeptide enrichment, and pY peptide enrichment prior to drying by vacuum centrifugation.

### R2P2 for global phosphopeptide enrichment

Enrichment was performed on the KingFisher flex as previously described^29^. Specifically,

1. Each 250 µg aliquot of dried peptides was resuspended in 900 µL 80% ACN, 0.1% TFA.
2. Precipitates were removed by centrifugation for 10 min at 21,000 × g and 4°C.
3. Clarified peptides were added to a deep 96-well plate leaving behind 50 µL to avoid transfer of precipitate.
4. Washes were prepared by filling three shallow 96-well plates with 150 µL of 80% ACN, 0.1% TFA.
5. The bead plate was prepared by adding 70 µL of 5% Fe^3+^ NTA magnetic beads in 80% ACN, 0.1% TFA.
6. A tip comb was nested in a shallow well plate.
7. The R2P2 global phosphopeptide enrichment protocol (Expanded View Content) was initiated on the KingFisher Flex.

a. At about 11 minutes remaining in the protocol, the elution plate was prepared by adding 75 µL of 2.5% ammonia, 50% ACN to wells and added during the programmed pause.
b. During the KingFisher method, two layers of C8 extraction disks were nested in a P200 tip and equilibrated with the following solutions using centrifugation cycles of 2 min at 300 x g:

i. 60 µL of 100% MeOH
ii. 60 µL of 100% ACN
iii. 60 µL of 70% ACN containing 0.25% AA
8. Eluates were acidified with 45 µL of 10% FA, 75% ACN immediately upon method completion.
9. Eluates were filtered through two layers of C8 material equilibrated as described above. Eluates were collected in collected in MS vials along with a subsequent wash of 60 µL 70% ACN, 0.25% AA.
10. Filtered eluates were dried by vacuum centrifugation and stored at -20°C until MS measurement.

### R2HaPpY with R2P2 clean-up

#### Preparation of peptide binding plate

1. Dried peptides ranging in amounts from 0.25 mg to 4 mg according to experiment, were resuspended in 900 µL of 50 mM Tris, 50 mM NaCl, pH 8.0 by vortexing and sonication.
2. Precipitate was pelleted by centrifugation ≥ 10,000 x g for 10 - 15 min at 4°C.
3. Clarified peptide solution was added to deep well KingFisher ‘Binding Plate’ leaving behind 50 µL to avoid adding precipitate.
4. Halo-TEV-sSrc beads were equilibrated into fresh 50 mM Tris, 50 mM NaCl, pH 8.0 by washing twice and resuspended at a 20% bead slurry concentration. A total of 40 µL of bead slurry was mixed with each peptide sample.

#### Preparation of tip comb, wash and elution plates

5. Tip plate: a shallow well plate held a deep well tip comb.
6. Wash plates:

a. Washes 1 – 3: three deep well plates received 850 µL of 50 mM Tris, 50 mM NaCl, pH 8.0 per well.
b. Wash 4: one deep well plate received 850 µL of HPLC-grade water.
7. Elution plates:

a. Elution plate 1: a shallow well elution plate received 100 µL of 0.5% TFA per well.
b. Elution plate 2 was prepared and added during the last 15 minutes of the method. For this, a shallow well elution plate received 120 µL of 1% TFA, 60% ACN per well.

#### KingFisher method

8. The R2HaPpY KingFisher Flex BindIt protocol (Expanded View Content) was started with all plates except elution 2.
9. Elution 2 was prepared and added at the pause at ∼15 min left in the protocol.
10. Upon completion of enrichment, elution 2 was combined into elution 1.

a. Pooled eluates were transferred to Eppendorf tubes and diluted to reach 75% ACN in a total volume of 900 µL.
b. Any precipitates or residual beads were pelleted by centrifugation for 4 min at 16,000 × *g*.
c. Supernatant was transferred to a deep well plate for additional clean-up using the R2P2 global phosphopeptide enrichment protocol described above with the following two differences.

i. The deep well plate containing the clarified pY peptide eluate at 75% ACN prepared in the above step served as the peptide binding plate.
ii. The bead plate contained 80 µL of 5% Fe-NTA IMAC bead slurry in 80% ACN, 0.1% TFA.
iii. All other steps described above were the same for pY peptide clean-up.
11. Dried pY peptide samples were resuspended in 12 µL of 4% ACN, 3% FA, 93% water.

### Mass spectrometry measurement

All measurements were performed on an Exploris 480 using either an Easy nLC 1200 or Easy nLC II. The EGF stimulation experiment used an Easy nLC II while all other experiments used an Easy nLC 1200. Peptides were loaded onto a 100 µm ID x 3 cm silica trap column packed with Reprosil C18 3 µm beads (Dr. Maisch GmbH). Peptides were separated along a 100 µm ID x 30 cm analytical column packed Reprosil C18 3 µm beads for 2 cm in the tip followed by 28 cm of Reprosil C18 1.9 µm beads (Dr. Maisch GmbH). The analytical column is housed in a column heater, set to 50°C and voltage was applied between the pre- and analytical columns.

### Tyrosine phosphoproteome measurements

Peptides were eluted using a gradient of 6.4% to 32% ACN in 66 min followed by 32% to 48% ACN in 7 min followed by 76% ACN for 6 min and 2.4% ACN for 11 min using a flow rate of 300-400 nL/min where solvent A is HPLC-grade 0.1% formic acid and solvent B is HPLC-grade 80% ACN, 0.1% formic acid.

Full MS scans were acquired from 375 to 1500 m/z at 120,000 resolution with a fill target of 3e6 ions and maximum injection time set to automatic. The most abundant ions on the full MS scan were selected for fragmentation using a 1.6 m/z precursor isolation window and beam-type collisional-activation dissociation (HCD) with 30% normalized collision energy for a cycle time of 3 s. Precursor intensity threshold was set at 2.5e4, and only precursors with charge states determined to be 2 – 6 were fragmented. MS/MS spectra were collected at 45,000 resolution with a fill target of 2e5 ions and maximum injection time of 86 ms.

For high phosphotyrosine containing samples such as pervanadate treated HeLa and Yeast vSrc, precursors were dynamically excluded from selection for 45 sec. For low phosphotyrosine containing samples, such as untreated or EGF-stimulated HeLa cells, dynamic exclusion settings were programmed to allow precursor fragmentation up to twice within 20 s prior to exclusion for 20s.

### Global phosphoproteome measurements

Analysis of global phosphoproteome was same as tyrosine phosphoproteome except with different LC gradient and dynamic exclusion settings. Specifically, phosphopeptides were separated by increasing amounts of ACN in 0.1% FA over 73 min gradient specifically ranging from 3.2% to 6.4% ACN in 3 minutes, 6.4% to 25.6% ACN in 63 minutes, 25.6% to 48% ACN in 7 minutes followed by 76% ACN for 6 min and 2.4% ACN for 11 min using a flow rate of 400 nL/min. Fragmented precursors were dynamically excluded from selection for 45 s.

### Total proteome measurements

Dried 10 µg aliquots of tryptic peptides representing the total proteome of HeLa cells in the EGF stimulation experiment were resuspended to 0.5 µg/µL in 3% ACN, 4% FA for measurement of 0.5 µg by data dependent acquisition. Peptides were separated by a 43-min gradient ranging from 6.4% to 36% acetonitrile in 0.1% formic acid using a flow rate of 350 nL/min. Full MS scans were acquired from 350 to 1500 m/z at 120,000 resolution with a fill target of 3e6 ions and maximum injection time set to automatic. The 15 most abundant ions on the full MS scan were selected for fragmentation using 2 m/z precursor isolation windows and beam-type collisional-activation dissociation (HCD) with 30% normalized collision energy. MS/MS spectra were collected at 15,000 resolution with the standard fill target and maximum injection time fixed to 40 ms. Fragmented precursors were dynamically excluded from selection for 30 s.

## Data analysis

### All datasets

DDA-MS/MS spectra were searched with Comet (release 2019.01.2)^137^ against the human proteome downloaded from Uniprot on March 21^st^, 2024 including only reviewed entries and excluding isoforms (20,418 entries). The yeast samples were searched against the *S. cerevisiae* proteome downloaded from Uniprot on February 19^th^, 2021 with the vSrc kinase sequence appended. Peptides were searched with trypsin digestion allowing up to two missed cleavages. The precursor *m/z* tolerance was set to 20 ppm. The fragment *m/z* mass tolerance was set to 0.02 Da. Constant modification of cysteine carbamidomethylation (57.02146372118 Da) and variable modification of methionine oxidation (15.9949146202 Da, max 2), and N-terminal acetylation (42.01056468472, max 1) were used for all searches. An additional variable modification of serine, threonine, and tyrosine phosphorylation (79.966331 Da, max 3) was used for global and tyrosine-specific phosphopeptide samples. Search results were filtered to a 1% FDR at PSM level using Percolator^138^. Phosphorylation sites were localized using an in-house implementation of the Ascore algorithm^139^. Phosphorylation sites with an Ascore ≥ 13 (*P* ≤ 0.05) were considered confidently localized. Peptides were quantified using in-house implementation of the moFF algorithm^140^ to extract MS1 peak maximum intensities.

### Bioinformatic analyses

Bioinformatic analysis was performed using R (https://www.r-project.org/). Quantitative values were consolidated by keeping the maximum intensity PSM per precursor ion (peptide sequence with charge state). For total proteome analysis, precursor ions were summed to the peptide level or to protein level. For phosphorylation site level analysis, all peptides containing the confidently localized phosphosite were summed. For phosphosite isoform analysis, all peptides containing the same one to three co-occurring and confidently localized phosphosites were summed. Intensity distributions of peptides, proteins, phosphosites, or phosphoisoforms were median normalized across conditions for each experiment.

For all boxplots, the lower and upper hinges of the boxes correspond to the 25% and 75% percentile, and the bar in the box to the median. The upper and lower whiskers extend from the highest and lowest values, respectively, but no further than 1.5 times the IQR from the hinge.

All correlation calculations utilize Pearson’s method.

### Bioinformatic analysis of EGF signaling response

#### Completeness filtering

A completeness filter was applied prior to analysis of biological insights from EGF stimulation data. Specifically, pY sites were required to be observed in at least 3 out of 6 biological replicates in minimum of one out of five conditions (EGF treatments or untreated). For pY sites passing the completeness criteria, missing values were imputed by sampling a random normal distribution as described below. Imputation was applied only to tests of differential abundance and temporal response profiling for the EGF stimulation experiment (Figure 5).

### Imputation

Imputation was applied to missing replicates prior to differential abundance testing to capture pY sites that were completely undetected in one condition yet confidently measured in another. For sites missing in all biological replicates of a condition, log_2_-transformed intensities were imputed by sampling a random normal distribution centered at 10 with a standard deviation of 1. The intensity distribution was centered at 10 to represent intensities slightly below the limit of detection given the lowest observed log_2_-transformed intensity was 11.02 and the lowest 1% percentile of log_2_-transformed intensity was 15.02. A standard deviation of 1 was chosen to provide a slightly greater variation than the average observed standard deviation of 0.63 to avoid inflated significance values. Additionally, a wider variation was chosen for the fully missing values due to higher variability inherent to lower abundance measurements. When pY sites were not detected in all biological replicates of one condition but were detected in at least 50% of replicates in a different condition, this imputation approach assigned values between 8 and 12 95% of the time. For sites detected in at least one replication of a condition, missing observations were imputed by sampling a randomized normal distribution, centered at the average intensity of measured replicates, with a standard deviation set to global average standard deviation of 0.63.

Missing observations were also imputed prior to temporal response profiling by fuzzy c-means clustering (Figure 3C and 5A, B), except without sampling from a randomized normal distribution. Instead, missing observations from all replicates of a condition received a log_2_-transformed intensity of 10, while missing observations from five or less replicates received a log_2_-transformed intensity set to the average intensity of the measured replicates. By contrast, plotting of measured site intensities, such as in Figure 4, 5C, and 6, only showed measured observations, excluding any imputed intensities.

### Differential abundance testing

The calculation of individual *p*-values used Welch’s two-sided *t*-test of biological replicates, using the untreated condition as control and applying Benjamini-Hochberg procedure for multiple testing correction. Phosphosites were considered differentially abundant fold change was greater than 1 or less than -1 and if adjusted *p*-value less than 0.05. The protti R package version 0.8.0^141^ was used for calculation of differential abundance. Functions for correlation analysis and coefficient of variation were based on Protti functions with added aesthetic modifications.

### Database Annotation analysis

For comparison of EGF-regulated pY sites in this study to databased EGF response signatures, two databases were downloaded. The first database was retrieved from the Site Search tool at PhosphoSitePlus.org^71^ on December 19^th^, 2024 specifying treatment as EGF. The second database was downloaded from PTMSigDB v2.0.0^104^ on September 3, 2024 for the semi-automatically curated EGFR1 perturbation dataset. Regulated pY sites identified in this study were joined to both databases by uniprot id, gene name and pY site location within full protein.

### Motif enrichment analyses

Sequence logos presented in Figure 4 were created with the R package ggseqlogo^142^ using as input the unique set of 15 amino acids flanking the regulated pY site. Logos were expressed in Bits. The total height of the letter stack at each position indicates the relative conservation at each position while the height of individual letters indicates the relative frequency of each amino acid within the position.

### Clustering with temporal response profiling

Clustering was performed by fuzzy c-means soft clustering, specifically using the fcm function from the e1071 R package (https://rdocumentation.org/packages/e1071/versions/1.7-16). Only pY sites measured in at least 50% of replicates of one timepoint and determined to be significantly regulated with an absolute log_2_(fold-change) greater than one and adjusted-pvalue less than 0.05 were included in clustering analysis. Imputation of missing replicates was performed as described above. pY site intensities were scaled prior to clustering. Clusters were assigned according to the maximum correlation to the overall cluster trend as determined by the fcm method. The total number of clusters representing the distinct temporal response profiles were chosen based on k-means grouping method described subsequently, coupled with manual inspection for the minimal number of clusters showing distinct trends. The cluster determination by k-means was performed according to the elbow method by choosing number of clusters at which the rate of increase in the amount of variation explained by adding another cluster drastically slowed (according to sum of squares divided by total sum of squares ratio calculated by the ‘kmeans’ function in the stats package in R (https://rdocumentation.org/packages/stats/versions/3.6.2)). We settled on four clusters because the amount of variation explained by adding a fifth cluster dropped from a 9% increase to a 3% increase and visual inspection determined each cluster described a distinct intensity profile.

### Gene ontology profiling

Gene ontology enrichment analysis was performed for Figure 5D and Figure EV13A-B. Unique proteins containing at least one regulated pY site were subset according to cluster and analyzed with gprofiler2 using the R package. To avoid overly broad functional enrichments, gene ontology terms were filtered for only terms containing 1000 or less children terms. Significant enrichment terms were then manually curated to avoid semantic redundancies.

Visualization of cluster distributions for pY sites on proteins within selected functional annotations was performed for Figure EV13C. Protein functional annotations were selected for direct comparison to Olsen *et al*. *Cell* 2006^85^. Functional annotations were assigned by mathcing of uniprot keyword ids to proteins with regulated pY sites. Specifically, KW-0418 indicated ‘kinases’; KW-0343 or term “guanine nucleotide exchange factor” in protein names indicated ‘GEFs & GAPs’; KW-0805 indicated ‘transcriptional regulators’; KW-0010 or KW-0678 indicated transcription factors; KW-0832 or KW-0833 indicateed ubiquitin ligases; KW-0009 indicated ‘actin binding’; KW-0694 indicated ‘RNA binding’; remaining unclassified proteins were labelled as ‘other’. Distributions of clusters within each protein function category were calculated based on counts of indiviudal pY sites.

## Data Availability

The mass spectrometry proteomics data have been deposited to the ProteomeXchange Consortium via the PRIDE partner repository^143,144^ with the dataset identifier PXD062515 and 10.6019/PXD062515

## Author Contributions

A.C. experimentation, design, analysis, writing. R. R-M. ideation, design, analysis, writing. M.B. experimentation, design, analysis, writing. S.M. experimentation. J.V. ideation, design, analysis, writing, funding.

## Disclosure and competing interests statement

The authors declare no competing interests.

## Acknowledgements

We thank Ariadna Llovet and Julian Ramos for providing support with in-house proteomics software, Alex Hogrebe for illustrations of instrumentation used in figures, Christian Landry (U. Laval) for gifting the yeast vSrc strain, and Robert Lawrence, Mario Leutert and Rachael Hu for helpful discussions. Research reported in this publication was primarily supported by the National Institute of General Medical Sciences of the National Institutes of Health under award number R35GM152061. The work and personnel involved in the project were additionally supported by NIH grants R35GM119536 from the National Institute of General Medicine and R01AG056359 from the National Institute of Aging; and by Human Frontiers Science Program grant RGP0034/2018. AC was supported by the National Human Genome Research Institute of the National Institutes of Health NIH under training grant T32HG000035. MDB was supported by a Canadian Institutes of Health Research Postdoctoral Fellowship (MFE-193932). The content is solely the responsibility of the authors and does not necessarily represent the official views of the National Institutes of Health or other funding agencies.

## Supplementary Material

1. Expanded View Content
2. Dataset EV1 – Lists of phosphosites and phosphopeptides
3. Dataset EV2 - List of novel pY sites
4. Dataset EV3 - List of node depth assignments for EGF pathway analysis

## References

1. Leutert, M., Barente, A. S., Fukuda, N. K., Rodriguez-Mias, R. A. & Villén, J. The regulatory landscape of the yeast phosphoproteome. bioRxiv 2022.10.23.513432 (2022) doi:10.1101/2022.10.23.513432.

2. Friedman, A. & Perrimon, N. Genetic screening for signal transduction in the era of network biology. Cell 128, 225–231 (2007).

3. Breitkreutz, A. et al. A global protein kinase and phosphatase interaction network in yeast. Science 328, 1043–1046 (2010).

4. Bodenmiller, B. et al. Phosphoproteomic analysis reveals interconnected system-wide responses to perturbations of kinases and phosphatases in yeast. Sci. Signal. 3, rs4 (2010).

5. Tan, C. S. H. et al. Comparative analysis reveals conserved protein phosphorylation networks implicated in multiple diseases. Sci. Signal. 2, ra39 (2009).

6. Swaney, D. L. et al. Global analysis of phosphorylation and ubiquitylation cross-talk in protein degradation. Nat. Methods 10, 676–682 (2013).

7. Ochoa, D. et al. The functional landscape of the human phosphoproteome. Nat. Biotechnol. 38, 365– 373 (2020).

8. Hunter, T. Tyrosine phosphorylation: thirty years and counting. Curr. Opin. Cell Biol. 21, 140–146 (2009).

9. Blume-Jensen, P. & Hunter, T. Oncogenic kinase signalling. Nature 411, 355–365 (2001).

10. Cohen, P., Cross, D. & Jänne, P. A. Kinase drug discovery 20 years after imatinib. Nat. Rev. Drug Discov. (2022) doi:10.1038/s41573-022-00418-2.

11. Rikova, K. et al. Global survey of phosphotyrosine signaling identifies oncogenic kinases in lung cancer. Cell 131, 1190–1203 (2007).

12. Guo, A. et al. Signaling networks assembled by oncogenic EGFR and c-Met. Proc. Natl. Acad. Sci. U. S. A. 105, 692–697 (2008).

13. Reddy, R. J. et al. Early signaling dynamics of the epidermal growth factor receptor. Proc. Natl. Acad. Sci. U. S. A. 113, 3114–3119 (2016).

14. Oyama, M. et al. Temporal perturbation of tyrosine phosphoproteome dynamics reveals the system-wide regulatory networks. Mol. Cell. Proteomics 8, 226–231 (2009).

15. Sharma, K. et al. Ultradeep human phosphoproteome reveals a distinct regulatory nature of Tyr and Ser/Thr-based signaling. Cell Rep. 8, 1583–1594 (2014).

16. Lundby, A. et al. Oncogenic Mutations Rewire Signaling Pathways by Switching Protein Recruitment to Phosphotyrosine Sites. Cell 179, 543–560.e26 (2019).

17. Francavilla, C. et al. Multilayered proteomics reveals molecular switches dictating ligand-dependent EGFR trafficking. Nat. Struct. Mol. Biol. 23, 608–618 (2016).

18. Sebé-Pedrós, A. et al. High-Throughput Proteomics Reveals the Unicellular Roots of Animal Phosphosignaling and Cell Differentiation. Dev. Cell 39, 186–197 (2016).

19. Yoshida, T. et al. Tyrosine phosphoproteomics identifies both codrivers and cotargeting strategies for T790M-related EGFR-TKI resistance in non-small cell lung cancer. Clin. Cancer Res. 20, 4059–4074 (2014).

20. Abe, Y. et al. Deep phospho- and phosphotyrosine proteomics identified active kinases and phosphorylation networks in colorectal cancer cell lines resistant to cetuximab. Sci. Rep. 7, 10463 (2017).

21. Invergo, B. M. & Beltrao, P. Reconstructing phosphorylation signalling networks from quantitative phosphoproteomic data. Essays Biochem. 62, 525–534 (2018).

22. Block, H. et al. Immobilized-metal affinity chromatography (IMAC): a review. Methods Enzymol. 463, 439–473 (2009).

23. Andersson, L. & Porath, J. Isolation of phosphoproteins by immobilized metal (Fe3+) affinity chromatography. Anal. Biochem. 154, 250–254 (1986).

24. Muszyńska, G., Andersson, L. & Porath, J. Selective adsorption of phosphoproteins on gel-immobilized ferric chelate. Biochemistry 25, 6850–6853 (1986).

25. Posewitz, M. C. & Tempst, P. Immobilized gallium(III) affinity chromatography of phosphopeptides. Anal. Chem. 71, 2883–2892 (1999).

26. Thingholm, T. E. & Larsen, M. R. The use of titanium dioxide for selective enrichment of phosphorylated peptides. Methods Mol. Biol. 1355, 135–146 (2016).

27. Thingholm, T. E. & Larsen, M. R. Phosphopeptide enrichment by immobilized metal affinity chromatography. Methods Mol. Biol. 1355, 123–133 (2016).

28. Tape, C. J. et al. Reproducible automated phosphopeptide enrichment using magnetic TiO2 and Ti-IMAC. Anal. Chem. 86, 10296–10302 (2014).

29. Leutert, M., Rodríguez-Mias, R. A., Fukuda, N. K. & Villén, J. R2-P2 rapid-robotic phosphoproteomics enables multidimensional cell signaling studies. Mol. Syst. Biol. 15, (12/2019).

30. Bergström Lind, S., et al. Immunoaffinity enrichments followed by mass spectrometric detection for studying global protein tyrosine phosphorylation. J. Proteome Res. 7, 2897–2910 (2008).

31. Bergström Lind, S., et al. Toward a comprehensive characterization of the phosphotyrosine proteome. Cell. Signal. 23, 1387–1395 (2011).

32. Artemenko, K. A. et al. Optimization of immunoaffinity enrichment and detection: toward a comprehensive characterization of the phosphotyrosine proteome of K562 cells by liquid chromatography-mass spectrometry. Analyst 136, 1971–1978 (2011).

33. Boersema, P. J. et al. In-depth qualitative and quantitative profiling of tyrosine phosphorylation using a combination of phosphopeptide immunoaffinity purification and stable isotope dimethyl labeling. Mol. Cell. Proteomics 9, 84–99 (2010).

34. Kettenbach, A. N. & Gerber, S. A. Rapid and Reproducible Single-Stage Phosphopeptide Enrichment of Complex Peptide Mixtures: Application to General and Phosphotyrosine-Specific Phosphoproteomics Experiments. Anal. Chem. 83, 7635–7644 (2011).

35. van der Mijn, J. C. et al. Evaluation of different phospho-tyrosine antibodies for label-free phosphoproteomics. J. Proteomics 127, 259–263 (2015).

36. Abe, Y., Nagano, M., Tada, A., Adachi, J. & Tomonaga, T. Deep Phosphotyrosine Proteomics by Optimization of Phosphotyrosine Enrichment and MS/MS Parameters. J. Proteome Res. 16, 1077–1086 (2017).

37. Steen, H., Kuster, B., Fernandez, M., Pandey, A. & Mann, M. Tyrosine phosphorylation mapping of the epidermal growth factor receptor signaling pathway. J. Biol. Chem. 277, 1031–1039 (2002).

38. Rush, J. et al. Immunoaffinity profiling of tyrosine phosphorylation in cancer cells. Nat. Biotechnol. 23, 94–101 (2005).

39. Blagoev, B., Ong, S.-E., Kratchmarova, I. & Mann, M. Temporal analysis of phosphotyrosine-dependent signaling networks by quantitative proteomics. Nat. Biotechnol. 22, 1139–1145 (2004).

40. Bian, Y. et al. Ultra-deep tyrosine phosphoproteomics enabled by a phosphotyrosine superbinder. Nat. Chem. Biol. 12, 959–966 (11/2016).

41. Humphrey, S. J., Karayel, O., James, D. E. & Mann, M. High-throughput and high-sensitivity phosphoproteomics with the EasyPhos platform. Nat. Protoc. 13, 1897–1916 (2018).

42. Humphrey, S. J., Azimifar, S. B. & Mann, M. High-throughput phosphoproteomics reveals in vivo insulin signaling dynamics. Nat. Biotechnol. 33, 990–995 (2015).

43. Klaeger, S. et al. The target landscape of clinical kinase drugs. Science 358, (2017).

44. Zecha, J. et al. Decrypting drug actions and protein modifications by dose- and time-resolved proteomics. Science 380, 93–101 (2023).

45. Fazakerley, D. J. et al. Phosphoproteomics reveals rewiring of the insulin signaling network and multi-nodal defects in insulin resistance. Nat. Commun. 14, 923 (2023).

46. Martinez-Val, A. et al. Proteomics of colorectal tumors identifies the role of CAVIN1 in tumor relapse. Mol. Syst. Biol. (2025) doi:10.1038/s44320-025-00102-8.

47. Martinez-Val, A. et al. Spatial-proteomics reveals phospho-signaling dynamics at subcellular resolution. Nat. Commun. 12, 7113 (2021).

48. Ladbury, J. E. et al. Measurement of the binding of tyrosyl phosphopeptides to SH2 domains: a reappraisal. Proc. Natl. Acad. Sci. U. S. A. 92, 3199–3203 (1995).

49. Ladbury, J. E. & Arold, S. T. Energetics of Src homology domain interactions in receptor tyrosine kinase-mediated signaling. Methods Enzymol. 488, 147–183 (2011).

50. Jones, R. B., Gordus, A., Krall, J. A. & MacBeath, G. A quantitative protein interaction network for the ErbB receptors using protein microarrays. Nature 439, 168–174 (2005).

51. Kaneko, T. et al. Loops govern SH2 domain specificity by controlling access to binding pockets. Sci. Signal. 3, ra34 (2010).

52. Kaneko, T., Joshi, R., Feller, S. M. & Li, S. S. Phosphotyrosine recognition domains: the typical, the atypical and the versatile. Cell Commun. Signal. 10, 32 (2012).

53. Machida, K. et al. High-throughput phosphotyrosine profiling using SH2 domains. Mol. Cell 26, 899– 915 (2007).

54. Tinti, M. et al. The SH2 domain interaction landscape. Cell Rep. 3, 1293–1305 (2013).

55. Kaneko, T. et al. Superbinder SH2 Domains Act as Antagonists of Cell Signaling. Sci. Signal. 5, (2012).

56. Martyn, G. D. et al. Engineered SH2 Domains for Targeted Phosphoproteomics. ACS Chem. Biol. (2022) doi:10.1021/acschembio.2c00051.

57. Li, Y., Wang, Y., Dong, M., Zou, H. & Ye, M. Sensitive Approaches for the Assay of the Global Protein Tyrosine Phosphorylation in Complex Samples Using a Mutated SH2 Domain. Anal. Chem. 89, 2304–2311 (2017).

58. Li, S. et al. Revisiting the phosphotyrosine binding pocket of Fyn SH2 domain led to the identification of novel SH2 superbinders. Protein Sci. 30, 558–570 (2021).

59. Li, A., Voleti, R., Lee, M., Gagoski, D. & Shah, N. H. High-throughput profiling of sequence recognition by tyrosine kinases and SH2 domains using bacterial peptide display. bioRxiv 2022.08.01.502334 (2022) doi:10.1101/2022.08.01.502334.

60. Yao, Y. et al. SH2 Superbinder Modified Monolithic Capillary Column for the Sensitive Analysis of Protein Tyrosine Phosphorylation. J. Proteome Res. 17, 243–251 (2018).

61. Yao, Y. et al. One-Step SH2 Superbinder-Based Approach for Sensitive Analysis of Tyrosine Phosphoproteome. J. Proteome Res. 18, 1870–1879 (2019).

62. Tong, J. et al. Protein-phosphotyrosine proteome profiling by superbinder-SH2 domain affinity purification mass spectrometry, sSH2-AP-MS. Proteomics **17**, (2017).

63. Chua, X. Y. et al. Tandem Mass Tag Approach Utilizing Pervanadate BOOST Channels Delivers Deeper Quantitative Characterization of the Tyrosine Phosphoproteome. Mol. Cell. Proteomics 19, 730–743 (04/2020).

64. Dong, M. et al. Sensitive, Robust, and Cost-Effective Approach for Tyrosine Phosphoproteome Analysis. Anal. Chem. 89, 9307–9314 (2017).

65. Chang, A., Leutert, M., Rodriguez-Mias, R. A. & Villén, J. Automated Enrichment of Phosphotyrosine Peptides for High-Throughput Proteomics. J. Proteome Res. 22, 1868–1880 (2023).

66. Kong, Q. et al. Integrated and High-Throughput Approach for Sensitive Analysis of Tyrosine Phosphoproteome. Anal. Chem. 94, 13728–13736 (2022).

67. Ke, A.-Q. et al. Development of novel affinity reagents for detecting protein tyrosine phosphorylation based on superbinder SH2 domain in tumor cells. Anal. Chim. Acta 1032, 138–146 (2018).

68. Chua, X. Y. & Salomon, A. Ovalbumin antigen-specific activation of human T cell receptor closely resembles soluble antibody stimulation as revealed by BOOST phosphotyrosine proteomics. J. Proteome Res. 20, 3330–3344 (2021).

69. Callahan, A. et al. Deep phosphotyrosine characterisation of primary murine T cells using broad spectrum optimisation of selective triggering. Proteomics e2400106 (2024).

70. Los, G. V. et al. HaloTag: a novel protein labeling technology for cell imaging and protein analysis. ACS Chem. Biol. 3, 373–382 (2008).

71. Hornbeck, P. V. et al. PhosphoSitePlus: a comprehensive resource for investigating the structure and function of experimentally determined post-translational modifications in man and mouse. Nucleic Acids Res. 40, D261–70 (2012).

72. Hornbeck, P. V. et al. PhosphoSitePlus, 2014: mutations, PTMs and recalibrations. Nucleic Acids Res. 43, D512–20 (2015).

73. N Peterson, S. & Kwon, K. The HaloTag: Improving soluble expression and applications in protein functional analysis. Curr. Chem. Genomics 6, 8–17 (2012).

74. Jayavelu, A. K., et al. EasyAb: A high-throughput workflow for antibody-based PTM peptide enrichment method coupled to mass spectrometry. bioRxiv 2024.10.29.620939 (2024) doi:10.1101/2024.10.29.620939.

75. Bradley, D. et al. The fitness cost of spurious phosphorylation. EMBO J. 43, 4720–4751 (2024).

76. Srivastava, A. K. & St-Louis, J. Smooth muscle contractility and protein tyrosine phosphorylation. Mol. Cell. Biochem. 176, 47–51 (1997).

77. Cohen, P. The origins of protein phosphorylation. Nat. Cell Biol. 4, E127–30 (2002).

78. Cohen, P. Protein kinases--the major drug targets of the twenty-first century? Nat. Rev. Drug Discov. 1, 309–315 (2002).

79. Rix, U. & Superti-Furga, G. Target profiling of small molecules by chemical proteomics. Nat. Chem. Biol. 5, 616–624 (2009).

80. Hu, C. et al. Anti-EGFR therapy can overcome acquired resistance to the third-generation ALK-tyrosine kinase inhibitor lorlatinib mediated by activation of EGFR. Acta Pharmacol. Sin. 1–15 (2025).

81. Castoldi, R. et al. TetraMabs: simultaneous targeting of four oncogenic receptor tyrosine kinases for tumor growth inhibition in heterogeneous tumor cell populations. Protein Eng. Des. Sel. 29, 467–475 (2016).

82. Korkut, A. et al. Perturbation biology nominates upstream-downstream drug combinations in RAF inhibitor resistant melanoma cells. Elife 4, (2015).

83. Flobak, Å. et al. Discovery of drug synergies in gastric cancer cells predicted by logical modeling. PLoS Comput. Biol. 11, e1004426 (2015).

84. Beekhof, R. et al. INKA, an integrative data analysis pipeline for phosphoproteomic inference of active kinases. Mol. Syst. Biol. 15, e8250 (2019).

85. Olsen, J. V. et al. Global, in vivo, and site-specific phosphorylation dynamics in signaling networks. Cell 127, 635–648 (2006).

86. Wind, S., Schnell, D., Ebner, T., Freiwald, M. & Stopfer, P. Clinical Pharmacokinetics and Pharmacodynamics of Afatinib. Clin. Pharmacokinet. 56, 235–250 (2017).

87. Ocana, A. et al. HER3 overexpression and survival in solid tumors: a meta-analysis. J. Natl. Cancer Inst. 105, 266–273 (2013).

88. Li, D. et al. BIBW2992, an irreversible EGFR/HER2 inhibitor highly effective in preclinical lung cancer models. Oncogene 27, 4702–4711 (2008).

89. Solca, F. et al. Target binding properties and cellular activity of afatinib (BIBW 2992), an irreversible ErbB family blocker. J. Pharmacol. Exp. Ther. 343, 342–350 (2012).

90. Kohale, I. N. et al. Quantitative Analysis of Tyrosine Phosphorylation from FFPE Tissues Reveals Patient-Specific Signaling Networks. Cancer Res. 81, 3930–3941 (2021).

91. Harrison, P. T., Vyse, S. & Huang, P. H. Rare epidermal growth factor receptor (EGFR) mutations in non-small cell lung cancer. Semin. Cancer Biol. 61, 167–179 (2020).

92. Kong, Q. et al. Dynamic phosphotyrosine-dependent signaling profiling in living cells by two-dimensional proximity proteomics. J. Proteome Res. 21, 2727–2735 (2022).

93. Kholodenko, B. N., Demin, O. V., Moehren, G. & Hoek, J. B. Quantification of short term signaling by the epidermal growth factor receptor. J. Biol. Chem. 274, 30169–30181 (1999).

94. Offterdinger, M., Georget, V., Girod, A. & Bastiaens, P. I. Imaging phosphorylation dynamics of the epidermal growth factor receptor. Journal of Biological Chemistry 279, 36972–36981 (2004).

95. Liu, F. & Roth, R. A. Grb-IR: a SH2-domain-containing protein that binds to the insulin receptor and inhibits its function. Proc. Natl. Acad. Sci. U. S. A. 92, 10287–10291 (1995).

96. Dong, L. Q., Porter, S., Hu, D. & Liu, F. Inhibition of hGrb10 binding to the insulin receptor by functional domain-mediated oligomerization. J. Biol. Chem. 273, 17720–17725 (1998).

97. Nantel, A., Mohammad-Ali, K., Sherk, J., Posner, B. I. & Thomas, D. Y. Interaction of the Grb10 adapter protein with the Raf1 and MEK1 kinases. J. Biol. Chem. 273, 10475–10484 (1998).

98. Frantz, J. D., Giorgetti-Peraldi, S., Ottinger, E. A. & Shoelson, S. E. Human GRB-IRbeta/GRB10. Splice variants of an insulin and growth factor receptor-binding protein with PH and SH2 domains. J. Biol. Chem. **272**, 2659–2667 (1997).

99. Dong, L. Q. et al. Cloning, chromosome localization, expression, and characterization of an Src homology 2 and pleckstrin homology domain-containing insulin receptor binding protein hGrb10gamma. J. Biol. Chem. 272, 29104–29112 (1997).

100. Deng, Y., Zhang, M. & Riedel, H. Mitogenic roles of Gab1 and Grb10 as direct cellular partners in the regulation of MAP kinase signaling. J. Cell. Biochem. 105, 1172–1182 (2008).

101. Morrione, A., Valentinis, B., Resnicoff, M., Xu, S. q. & Baserga, R. The role of mGrb10alpha in insulin-like growth factor I-mediated growth. J. Biol. Chem. 272, 26382–26387 (1997).

102. Qin, Z. et al. Transketolase (TKT) activity and nuclear localization promote hepatocellular carcinoma in a metabolic and a non-metabolic manner. J. Exp. Clin. Cancer Res. 38, 154 (2019).

103. Leutert, M., Entwisle, S. W. & Villén, J. Decoding Post-Translational Modification Crosstalk With Proteomics. Mol. Cell. Proteomics 20, 100129 (2021).

104. Krug, K. et al. A Curated Resource for Phosphosite-specific Signature Analysis *[S]. Mol. Cell. Proteomics 18, 576–593 (2019).

105. El-Sibai, M. et al. Cdc42 is required for EGF-stimulated protrusion and motility in MTLn3 carcinoma cells. J. Cell Sci. 120, 3465–3474 (2007).

106. Surve, S., Watkins, S. C. & Sorkin, A. EGFR-RAS-MAPK signaling is confined to the plasma membrane and associated endorecycling protrusions. J. Cell Biol. 220, (2021).

107. Pinilla-Macua, I., Watkins, S. C. & Sorkin, A. Endocytosis separates EGF receptors from endogenous fluorescently labeled HRas and diminishes receptor signaling to MAP kinases in endosomes. Proc. Natl. Acad. Sci. U. S. A. 113, 2122–2127 (2016).

108. Kolberg, L., Raudvere, U., Kuzmin, I., Vilo, J. & Peterson, H. gprofiler2 -- an R package for gene list functional enrichment analysis and namespace conversion toolset g:Profiler. F1000Res. **9**, (2020).

109. Kolberg, L. et al. g:Profiler-interoperable web service for functional enrichment analysis and gene identifier mapping (2023 update). Nucleic Acids Res. 51, W207–W212 (2023).

110. Abd-Rabbo, D. & Michnick, S. W. Delineating functional principles of the bow tie structure of a kinase-phosphatase network in the budding yeast. BMC Syst. Biol. 11, 38 (2017).

111. Levy, E. D., Landry, C. R. & Michnick, S. W. Cell signaling. Signaling through cooperation. Science (New York, N.Y.) vol. 328 983–984 (2010).

112. Martens, M. et al. WikiPathways: connecting communities. Nucleic Acids Res. 49, D613–D621 (2021).

113. Ben-Levy, R. et al. Identification of novel phosphorylation sites required for activation of MAPKAP kinase-2. EMBO J. 14, 5920–5930 (1995).

114. Roberts, P. J. & Der, C. J. Targeting the Raf-MEK-ERK mitogen-activated protein kinase cascade for the treatment of cancer. Oncogene 26, 3291–3310 (2007).

115. Thines, L., Li, Z. & Sacks, D. B. IQGAP1 is a phosphotyrosine-regulated scaffold for SH2-containing proteins. Cells 12, 483 (2023).

116. Hedman, A. C. et al. Tyrosine phosphorylation of the scaffold protein IQGAP1 in the MET pathway alters function. J. Biol. Chem. 295, 18105–18121 (2020).

117. McNulty, D. E., Li, Z., White, C. D., Sacks, D. B. & Annan, R. S. MAPK scaffold IQGAP1 binds the EGF receptor and modulates its activation. J. Biol. Chem. 286, 15010–15021 (2011).

118. Park, T.-J. & Curran, T. Essential roles of Crk and CrkL in fibroblast structure and motility. Oncogene 33, 5121–5132 (2014).

119. Sriram, G. et al. Iterative tyrosine phosphorylation controls non-canonical domain utilization in Crk. Oncogene 34, 4260–4269 (2015).

120. Cheung, H. W. et al. Amplification of CRKL induces transformation and epidermal growth factor receptor inhibitor resistance in human non-small cell lung cancers. Cancer Discov. 1, 608–625 (2011).

121. Birge, R. B., Kalodimos, C., Inagaki, F. & Tanaka, S. Crk and CrkL adaptor proteins: networks for physiological and pathological signaling. Cell Commun. Signal. 7, 13 (2009).

122. Ren, R., Ye, Z. S. & Baltimore, D. Abl protein-tyrosine kinase selects the Crk adapter as a substrate using SH3-binding sites. Genes Dev. 8, 783–795 (1994).

123. Feller, S. M., Knudsen, B. & Hanafusa, H. c-Abl kinase regulates the protein binding activity of c-Crk. EMBO J. 13, 2341–2351 (1994).

124. Watanabe, T. et al. Crk adaptor protein-induced phosphorylation of Gab1 on tyrosine 307 via Src is important for organization of focal adhesions and enhanced cell migration. Cell Res. 19, 638–650 (2009).

125. Jankowski, W. et al. Domain organization differences explain Bcr-Abl’s preference for CrkL over CrkII. Nat. Chem. Biol. 8, 590–596 (2012).

126. Hashimoto, Y. et al. Phosphorylation of CrkII adaptor protein at tyrosine 221 by epidermal growth factor receptor. J. Biol. Chem. 273, 17186–17191 (1998).

127. Beitner-Johnson, D. & LeRoith, D. Insulin-like growth factor-I stimulates tyrosine phosphorylation of endogenous c-Crk. J. Biol. Chem. 270, 5187–5190 (1995).

128. Khwaja, A., Hallberg, B., Warne, P. H. & Downward, J. Networks of interaction of p120cbl and p130cas with Crk and Grb2 adaptor proteins. Oncogene 12, 2491–2498 (1996).

129. Ribon, V. & Saltiel, A. R. Nerve growth factor stimulates the tyrosine phosphorylation of endogenous Crk-II and augments its association with p130Cas in PC-12 cells. J. Biol. Chem. 271, 7375–7380 (1996).

130. Blakesley, V. A. et al. Sphingosine 1-phosphate stimulates tyrosine phosphorylation of Crk. J. Biol. Chem. 272, 16211–16215 (1997).

131. Moritz, A. et al. Akt-RSK-S6 kinase signaling networks activated by oncogenic receptor tyrosine kinases. Sci. Signal. 3, ra64 (2010).

132. Pesesse, X. et al. The Src homology 2 domain containing inositol 5-phosphatase SHIP2 is recruited to the epidermal growth factor (EGF) receptor and dephosphorylates phosphatidylinositol 3,4,5-trisphosphate in EGF-stimulated COS-7 cells. J. Biol. Chem. **276**, 28348–28355 (2001).

133. Huang, P. H. et al. Quantitative analysis of EGFRvIII cellular signaling networks reveals a combinatorial therapeutic strategy for glioblastoma. Proc. Natl. Acad. Sci. U. S. A. 104, 12867–12872 (2007).

134. Le Coq, J., López Navajas, P., Rodrigo Martin, B., Alfonso, C. & Lietha, D. A new layer of phosphoinositide-mediated allosteric regulation uncovered for SHIP2. FASEB J. 35, e21815 (2021).

135. Le Coq, J. et al. Structural basis for interdomain communication in SHIP2 providing high phosphatase activity. Elife 6, (2017).

136. Jørgensen, C. et al. Cell-specific information processing in segregating populations of Eph receptor ephrin-expressing cells. Science 326, 1502–1509 (2009).

137. Eng, J. K., Jahan, T. A. & Hoopmann, M. R. Comet: An open-source MS/MS sequence database search tool. Proteomics 13, 22–24 (2013).

138. Käll, L., Canterbury, J. D., Weston, J., Noble, W. S. & MacCoss, M. J. Semi-supervised learning for peptide identification from shotgun proteomics datasets. Nat. Methods 4, 923–925 (2007).

139. Beausoleil, S. A., Villén, J., Gerber, S. A., Rush, J. & Gygi, S. P. A probability-based approach for high-throughput protein phosphorylation analysis and site localization. Nat. Biotechnol. 24, 1285–1292 (2006).

140. Argentini, A. et al. moFF: a robust and automated approach to extract peptide ion intensities. Nat. Methods 13, 964–966 (2016).

141. Quast, J.-P., Schuster, D. & Picotti, P. protti: an R package for comprehensive data analysis of peptide- and protein-centric bottom-up proteomics data. Bioinform Adv 2, vbab041 (2022).

142. Wagih, O. ggseqlogo: a versatile R package for drawing sequence logos. Bioinformatics 33, 3645–3647 (2017).

143. Perez-Riverol, Y. et al. The PRIDE database at 20 years: 2025 update. Nucleic Acids Res. 53, D543–D553 (2025).

144. Perez-Riverol, Y. et al. The PRIDE database and related tools and resources in 2019: improving support for quantification data. Nucleic Acids Res. 47, D442–D450 (2019).

